# Tissue-specific apparent mtDNA heteroplasmy and its relationship with ageing and mtDNA gene expression

**DOI:** 10.1101/2024.12.11.627989

**Authors:** Simon Wengert, Xenofon Giannoulis, Peter Kreitmaier, Pauline Kautz, Leif S. Ludwig, Holger Prokisch, Francesco Paolo Casale, Matthias Heinig, Na Cai

**Affiliations:** Helmholtz Pioneer Campus, Helmholtz Munich, Neuherberg, Germany; Computational Health Centre, Helmholtz Munich, Neuherberg, Germany; School of Computation, Information and Technology, Technical University of Munich, Munich, Germany; Institute of Computational Biology, Helmholtz Munich, Neuherberg, Germany; Institute of Human Genetics, School of Health and Medicine, Technical University of Munich, Munich, Germany; Institute of Translational Genomics, Helmholtz Munich, Neuherberg, Germany; Technical University of Munich (TUM) and Klinikum Rechts der Isar, TUM School of Medicine and Health, 81675 Munich, Germany; Technical University of Munich (TUM), School of Medicine and Health, Graduate School of Experimental Medicine, Munich, 81675, Germany; Berlin Institute of Health at Charité-Universitätsmedizin Berlin, Berlin, Germany; Max-Delbrück-Center for Molecular Medicine in the Helmholtz Association (MDC), Berlin Institute for Medical Systems Biology (BIMSB), Berlin, Germany; Institute of Biotechnology, Technische Universität Berlin, Berlin, Germany; Institute of Neurogenomics, Helmholtz Munich, Munich, Germany; Institute of AI for Health, Helmholtz Munich, Munich, Germany

## Abstract

Heteroplasmy in the mitochondrial DNA (mtDNA) accumulates with age, but how much it occurs in tissues and how it affects function and physiology is not well-studied. Here we present a comprehensive tissue-specific map of mtDNA heteroplasmy and mtRNA modifications, and their relationships with age and mitochondrial gene expression. We propose a robust variant calling pipeline for mtDNA heteroplasmy and mtRNA modifications using bulk RNAseq data from 49 tissues in GTEx, and a calibrated phenotype association framework for both types of variants. We identify 109 associations between them and donor age, and 784 associations with tissue-specific mtDNA gene expression. Of these, 7 and 18 show cell-type specificity within tissue. In addition, we find 9 instances where these variations mediate donor age effects on mtDNA gene expression. Finally, we confirm previously identified relationships between mt-tRNA modifications and gene expression on their 5’, but read- and gene-level investigations reveal previously unmodeled complexities.

## Introduction

Mitochondrial DNA (mtDNA) heteroplasmy is a well-studied phenomenon in humans, where there is more than one allele at a single position in the mtDNA in a single individual, either due to the inheritance of more than one allele at this position through the maternal germline, or somatic mutations in the mtDNA that occur and accumulate during development and ageing. Numerous links have been established between mtDNA heteroplasmy and mitochondrial diseases^1^, various cancers^2,3^, cardiovascular disease^4^ and ageing^5–7^, corroborated by investigations in non-human mammals^8,9^. However, little is known about the mechanisms behind these links. For example, we have very little understanding of whether mtDNA heteroplasmy affects mtDNA gene expression, if these effects could be age-related, cell-type specific, and if they are potentially pathogenic. What roles mtDNA heteroplasmy plays in resulting in the phenotypic outcomes they are associated with remains to be identified.

To date most studies on mtDNA heteroplasmy are performed on whole-genome sequencing (WGS) data or targeted mtDNA sequencing data, particularly in tumour tissues aimed at identifying their variations and evolution among cancer cells^3,10–13^. One recent study focused their efforts instead on investigating mtDNA heteroplasmy turnover, drift and selection^14^ in hematopoietic stem cells and progenitor cells^15^, and another directly compared these dynamics using WGS data on clones of both non-cancerous and cancerous tissue^16^, generating a highly informative reference for future studies. The increasing exploration of the utility of single-cell ATAC sequencing (sc-ATACseq) and innovation of mtDNA-enriching single-cell RNA sequencing (sc-RNAseq)^17^ technologies has also enabled the development of analyses frameworks using mtDNA derived from scRNAseq or sc-ATACseq data for lineage tracing purposes^18–20^. Only a couple of studies^21,22^, as far as we are aware, assess levels of tissue-specific mtDNA heteroplasmy using either bulk or single-cell RNAseq data in primary, somatic tissues. These pioneering studies focused on establishing types of “apparent” heteroplasmic variations present in RNAseq data (**Figure 1A**), including those directly transcribed from mtDNA heteroplasmy, and those due to reverse transcription artefacts in cDNA library preparation that are in fact indicative of mtRNA methylations, on transcript positions 9 and 58 of mt-tRNAs, as well as transcript position 947 of *MT-RNR2* and position 834 of *MT-ND5*. Though the methylation processes at these sites have largely been identified^23–28^, their downstream consequences on mitochondrial RNA and protein abundance and function have not been well-interrogated.

**Figure 1:**
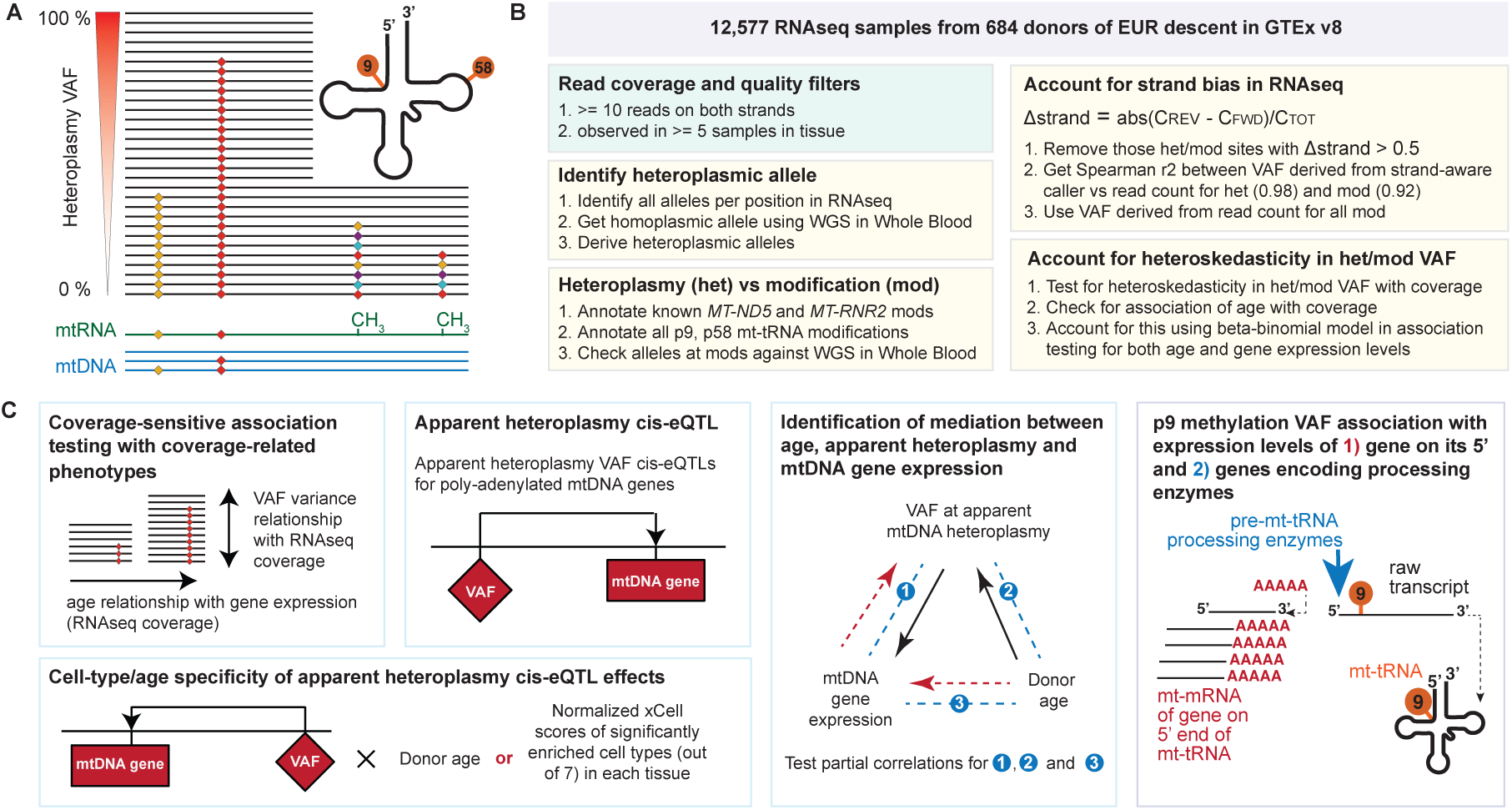
**A.** heteroplasmy levels detected in RNAseq alignment data can be reflecting either mtDNA mutations or post-transcriptional methylation events (-CH3) at specific sites in the mtRNA. These are jointly referred to as apparent heteroplasmy. **B.** Schematic showing the quality control pipeline we derive for analysing apparent mtDNA heteroplasmy in RNAseq samples from GTEx v8; the green box are related to sample/site level quality control that would be similar to mtDNA heteroplasmy identification from WGS data; the yellow boxes show operations specific to identifying and quantifying apparent mtDNA heteroplasmy from tissue-specific RNAseq data and that account for specific artefacts identified in RNAseq. **C.** Schematic showing our analysis pipeline using variant allele frequencies (VAF) at tissue-specific apparent mtDNA heteroplasmy. by whether the allele changes are transitions or transversions. **F.** Boxplot showing the levels of mtDNA heteroplasmy at the pathogenic mt.3243A>G variation in tissues it is identified in; boxes are coloured by tissue types. **G.** Scatterplot showing the number of mtDNA heteroplasmy of putatively somatic and inherited origins identified per tissue against the median mtDNA-CN levels shown in Rath SP et al.^48^; each dot represents a tissue; dots are coloured by the putative mtDNA heteroplasmy origin; lines visualizes the linear relationships between number of mtDNA heteroplasmy per tissue and median mtDNA-CN levels. **H.** Forest plot showing the enrichment odds ratio (OR) of mtRNA modifications mediated through different enzymes in CNS tissues, as opposed to other tissues; each dot represents all mtRNA modifications mediated through a single enzyme; dots are coloured by enzyme; error bars represent 95% confidence intervals (CI). For all boxplots, boxes show the interquartile range (IQR); horizontal lines in the boxes show median values; whiskers of boxes show 1.5xIQR.

In terms of associations between mtDNA heteroplasmy and phenotypes of interest, there have only been a few association studies to date, one with mtDNA copy number (mtDNA-CN)^29^, and another with the expression levels of mtDNA genes right beside the mtRNA modifications^22^. One reason for the lack of association studies between mtDNA heteroplasmy and phenotypes of interest may be the absence of well-calibrated statistical methods^30^, especially those that account for the effect of RNAseq coverage on variance in level mtDNA heteroplasmy. One recently proposed framework^31^ is mainly concerned with testing burdens of mtDNA heteroplasmy aggregated at the gene level, rather than the identification of effects on individual heteroplasmic positions. It is not targeted at identifying site-specific effects on gene expression regulation, mRNA degradation, and transcript processing, which we hope to assess. This calls for the development of a well-calibrated single-variant association testing framework for mtDNA heteroplasmy.

In this study we characterise, for the first time, apparent mtDNA heteroplasmic variations from bulk RNAseq data in 47 primary tissues and 2 cell lines in GTEx version 8 release^32^. As mtDNA heteroplasmy are tissue – and even cell-type – specific, and their levels vary as one ages, we postulate that assessing tissue-specific and age-dependent relationships between apparent heteroplasmy and gene expression levels presents the best way to investigate their effects on tissue function and physiology. We develop a pipeline to quantify apparent mtDNA heteroplasmy from RNAseq, accounting for sequencing artefacts characteristic of RNAseq data, and distinguishing between those arising from mtDNA mutations and those due to mtRNA modifications (**Figure 1B**). We further propose a two-step phenotype association testing framework for apparent mtDNA heteroplasmy, and demonstrate it is well-calibrated using simulations (**Figure 1C**). Applying this framework to RNAseq data in 49 tissues in GTEx v8, we identify 109 associations between apparent mtDNA heteroplasmy with donor age, and 784 associations with mtDNA gene expression levels, of which 7 and 18 show evidence of cell-type specificity. Notably, we find 9 instances with evidence of mtDNA heteroplasmy mediating donor age effects on mtDNA gene expression levels. Finally, we perform a direct investigation into whether mt-tRNA modifications on position 9 affect gene expression levels of their 5’ protein-coding gene as previously proposed^22^ using read-level data, and demonstrate that this relationship is likely more complex than previously established. In summary, our study provides the first comprehensive and tissue-specific description of the apparent mtDNA heteroplasmy, their relationship with donor age, and their regulatory effects on mitochondrial gene expression.

## Results

### Quantifying apparent mtDNA heteroplasmy from RNAseq data

To quantify tissue-specific apparent mtDNA heteroplasmy in RNA sequencing (RNAseq) data we perform heteroplasmy calling using mtDNA-sever^33^ in 12,577 RNAseq samples on 49 tissues obtained from 684 donors of European ancestry available in the GTEx v8 release^32^ (**Figure 1B, Supplementary Table S1, Methods**). In order to obtain heteroplasmy calls of high confidence we perform a series of stringent quality controls (**Supplementary Methods, Supplementary Table S1**). In particular, we identified the following noteworthy features in RNAseq data that may affect the accuracy of mtDNA heteroplasmic variant calls.

First, we find it is difficult to confidently infer the inherited and heteroplasmic alleles from RNAseq alone. Heteroplasmy levels can be different at the same position between different tissues (**Extended Data Figure 1**), consistent with our expectations of unequal segregation of inherited mtDNA heteroplasmy^16^, tissue-specific accumulation and drift of mtDNA mutations^14^, as well as tissue-specific levels of mtRNA modification^22^. Further, heteroplasmy can occur at homoplasmic variant sites, where different individuals in the GTEx dataset may have different inherited mtDNA alleles^34^. We therefore standardise the use of the major allele observed in WGS in Whole Blood as the inherited allele in all cases, and determine the variant allele frequency (VAFs, indicating levels of heteroplasmy) of the other alleles at the same positions in the same samples. This allows for VAFs to exceed 50% in some instances.

Second, we confirm previous observations of a sequencing artefact in RNAseq that occurs at mtRNA post-transcriptional modification sites^21,22^. These sites show the distinctive feature of having multiple heteroplasmic alleles (**Extended Data Figure 1**), consistent with the expectation of random base incorporation due to this sequencing artefact^21,22,35–38^, and do not show heteroplasmy in WGS (**Supplementary Figure S1**). We henceforth identify these sites as mtRNA modifications, distinguishing them from mtDNA heteroplasmy.

Third, we find pervasive strand bias (great difference in coverage between strands) in RNAseq data, especially at instances of heteroplasmy with low coverage, which resulted in apparent heteroplasmic positions in each sample with great discrepancies between VAF we obtained through mtDNA-server variant calling and those we obtain from raw read counts (**Extended Data Figure 1, Supplementary Figure S2, Supplementary Methods)**. These discrepancies are likely due to mtDNA-server’s maximum likelihood model being applied independently to each sequencing strand and demonstrate that mtDNA-server was not optimised for RNAseq data with pervasive strand bias^33^, mostly particularly in mtRNA modification sites (**Supplementary Figure S3**). We therefore used counts of heteroplasmic alleles across reads as VAFs at mtRNA modification sites with identified strand biases, while using VAF outputs from mtDNA-server at positions without strand bias (**Supplementary Methods**).

Finally, we compared VAFs identified in WGS and RNAseq at mtDNA heteroplasmy sites in the same samples in Whole Blood, and found that 29.26% are discrepant (**Extended Data Figure 1**, **Supplementary Figure S4**). We attribute this to heteroskedasticity in VAF estimates at different RNAseq coverage; the implications of this in association studies using VAF and coverage-derived phenotypes like gene expression levels have previously been mostly discussed in relation to allele specific expression (ASE)^39–41^. We therefore propose accounting for differences in coverage supporting the heteroplasmy calls in association testing, as we detail later in the paper.

### Landscape of common apparent mtDNA heteroplasmy across 49 tissues

We obtained a total of 142,392 apparent mtDNA heteroplasmy in 12,532 samples at 556 positions across 49 tissues (47 primary, 2 cell lines, **Methods**, **Supplementary Methods, Supplementary Tables S2**). All 556 mtDNA positions have apparent heteroplasmy in at least 5 samples in at least 1 of the 49 tissues, giving a range of 28 to 331 common apparent heteroplasmy positions per tissue (VAF: 0.006–0.982) that we take forward for all following investigations (**Figure 2A**). Overall, there is great variation between numbers of apparent heteroplasmy per individual across tissues (**Figure 2A**). Of the 556 apparent mtDNA heteroplasmy, 15 are previously reported as post-transcriptional m1A/G methylations: 12 and 1 of them are on positions 9 and 58 of mitochondrial tRNAs respectively, 1 is on mitochondrial rRNA *MT-RNR2*, and 1 is on the mRNA of gene *MT-ND5*. We refer to these apparent mtDNA heteroplasmy at these 15 positions collectively as mtRNA modifications (**Supplementary Table S3**). As expected, there are much greater numbers of bi- and tri-allelic heteroplasmy at mtRNA modifications than mtDNA heteroplasmy (**Figure 2B**).

**Figure 2:**
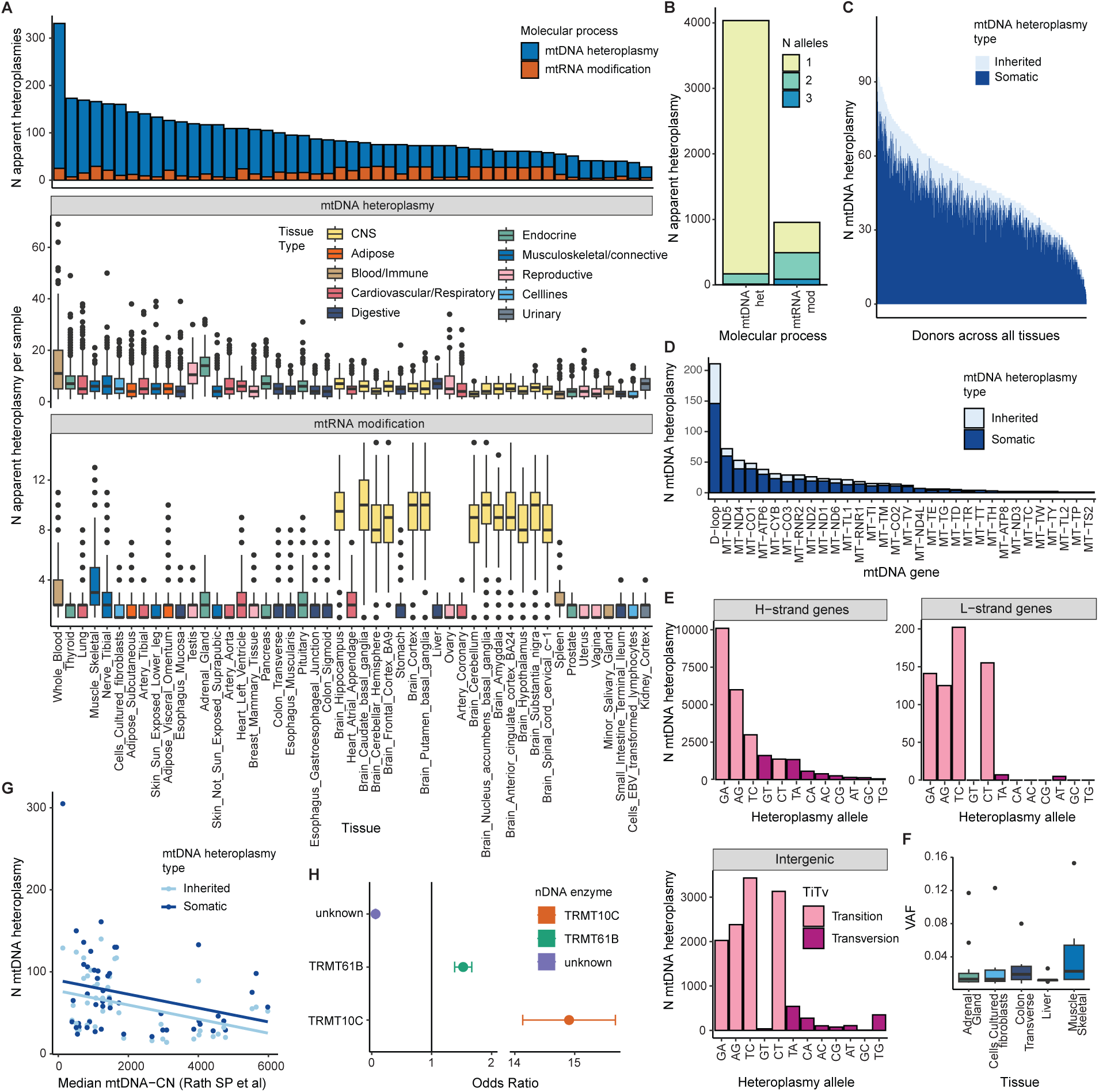
**A.** Upper: The total number of apparent heteroplasmy identified across all samples in each tissue called from RNAseq using the pipeline shown in Figure 1B; Lower: boxplots of number of apparent heteroplasmy sites identified in each donor across all tissues; each dot represents one sample; boxes are coloured by tissue types. **B.** Barplot showing the number of mtDNA heteroplasmy and mtRNA modifications with 1, 2 and 3 heteroplasmic alleles across all tissues; bars are coloured by the number of heteroplasmic alleles. **C.** Barplot showing the number of mtDNA heteroplasmy across tissues per donor that are putatively of somatic and inherited origins; bars are coloured by the putative mtDNA heteroplasmy origin. **D.** Barplot showing the number of mtDNA heteroplasmy of putatively inherited or somatic origin across all samples and tissues in each mtDNA gene; bars are coloured by the putative mtDNA heteroplasmy origin. **E.** Barplot showing the number of mtDNA heteroplasmy across all samples and tissues by heteroplasmy allele change; mtDNA heteroplasmy are grouped by their location in genes in the H-strand, genes in the L-strand, or intergenic regions; bars are coloured by whether the allele changes are transitions or transversions. **F.** Boxplot showing the levels of mtDNA heteroplasmy at the pathogenic mt.3243A>G variation in tissues it is identified in; boxes are coloured by tissue types. **G.** Scatterplot showing the number of mtDNA heteroplasmy of putatively somatic and inherited origins identified per tissue against the median mtDNA-CN levels shown in Rath SP et al.48; each dot represents a tissue; dots are coloured by the putative mtDNA heteroplasmy origin; lines visualizes the linear relationships between number of mtDNA heteroplasmy per tissue and median mtDNA-CN levels. **H.** Forest plot showing the enrichment odds ratio (OR) of mtRNA modifications mediated through different enzymes in CNS tissues, as opposed to other tissues; each dot represents all mtRNA modifications mediated through a single enzyme; dots are coloured by enzyme; error bars represent 95% confidence intervals (CI). For all boxplots, boxes show the interquartile range (IQR); horizontal lines in the boxes show median values; whiskers of boxes show 1.5xIQR.

For mtDNA heteroplasmy, we find the following: first, 0-48% of mtDNA heteroplasmic positions per donor are shared across 3 or more tissue types (**Figure 2C**), and therefore potentially inherited, while the remaining are more likely to have arisen due to somatic mutations in individual tissue lineages^16^. We find significant variation in the distribution of inherited or somatic mtDNA heteroplasmy between genes and tissues (**Figure 2D**, MANOVA P < 2×10^-16^ for both). At mtDNA heteroplasmy with putative inherited origin, we find higher variation in their VAFs among tissues in different tissue types than among those in the same tissue type (t-test P = 1.21×10^-4^). This suggests that uneven distribution of inherited mtDNA heteroplasmy in early cell divisions^42,43^ may result in higher variation in mtDNA heteroplasmy than selection and drift^44^.

Second, we find that 88.12% of the heteroplasmic alleles are transitions, consistent with previous findings^16^. In particular, G>A heteroplasmy make up the majority of mtDNA heteroplasmy of putative somatic origin identified on genes encoded by the heavy strand (H-strand, 40.6%), while all transitions more evenly contribute to mtDNA heteroplasmy found in genes encoded by the light strand (L-strand), even though the L-strand is depleted for purines, consistent with the hypothesis that most mtDNA heteroplasmy are results of errors on individual strands in mtDNA replication^16^ (**Figure 2E**). Third, consistent with previous reports^45–47^, mtDNA heteroplasmy at the pathogenic variant mt.3243A>G is only discovered in our dataset past VAF filters (**Methods**) in non-replicative tissues such as skeletal muscle and adrenal gland than replicative ones like liver and whole blood (where we do not observe the heteroplasmy at all, **Figure 2F**). Lastly, looking across tissues, we find that there are more mtDNA heteroplasmy of putative somatic origin in tissues with lower mtDNA copy number^48^ (mtDNA-CN, Pearson r = −0.41, P = 4.63×10^-3^, **Figure 2G**), consistent with greater effects of replication errors and drift at lower mtDNA-CN. In contrast, this effect is not significant for inherited mtDNA heteroplasmy, suggesting the effect of replication is potentially greater than that of drift (Pearson r = −0.27, P = 0.07, **Figure 2G**).

For mtRNA modifications, we find that they are present in higher numbers in tissues in the central nervous system (CNS, mean sites per donor = 15.4, se = 0.116) than all other tissues (mean sites per donor = 3.25, se = 0.0199). We therefore formally assess their CNS enrichment, taking into account the difference in sample size per tissue, as well as the non-random contribution of tissue samples by donors (**Supplementary Methods, Figure 2H**). To do this we grouped the mtRNA modifications into 3 sets, according to the enzymes that catalyse the modifications. We find that first set, containing the 12 m1A/G methylations on purine residues at mt-tRNA transcript position 9, catalysed by the nDNA encoded enzyme TRMT10C^23,24,26^, are significantly upregulated in CNS tissues than in other tissues (OR = 14.90, 95% CI = 14.10–15.70). The second set, containing just one m1A/G methylation at mt-tRNA transcript position 58, catalysed by the nDNA encoded enzyme TRMT61B^27^, is also significantly enriched in CNS (mean OR = 1.53, 95% CI = 1.38–1.67). Finally, the third set containing methylations at mt.2617 on *MT-RNR2* (transcript position 947) and mt.13710 on *MT-ND5* (transcript position 834), where the mediating enzyme is currently unknown^49^, is significantly depleted in the CNS (mean OR = 0.00635, 95% CI = 0.061– 0.067).

### Model selection for association testing in RNAseq derived heteroplasmy levels

Next, we set out to develop a framework for performing association testing between VAFs at apparent mtDNA heteroplasmy derived from RNAseq data and donor phenotypes, such as donor age, specifically taking into account any heteroskedasticity in VAFs at varying RNAseq coverage.

To quantify the problem of heteroskedasticity in association testing with VAFs, we first ask if apparent mtDNA heteroplasmy levels were significantly correlated with RNAseq coverage. We find that 2086 out of 4334 apparent mtDNA heteroplasmy positions are associated significantly with their site-specific RNAseq coverage (**Extended Data Figure 2, Supplementary Table S4**). Of the 4334 apparent mtDNA heteroplasmy positions, we find 70 apparent mtDNA heteroplasmy positions with significant heteroskedasticity in VAF due to differences in RNAseq coverage (**Supplementary Methods**, **Extended Data Figure 2, Supplementary Table S5**). We then ask whether RNAseq coverage at apparent mtDNA heteroplasmic positions is significantly associated with donor age, just like mtDNA-CN has previously been found to be^48,50^. We find that this is true at 830 out of 4,334 apparent mtDNA heteroplasmy positions across tissues (**Extended Data Figure 2, Supplementary Table S6**). Taken together, differences in RNAseq coverage underlying VAF estimates have the potential to induce spurious correlations between VAFs and donor phenotypes which may affect RNAseq coverage in turn, such as donor age.

Finally, we ask how this affects association testing using a canonical linear model (LM), and whether a Beta-Binomial (BB) regression model, commonly used for tackling similar issues in allele specific expression (ASE) testing^40,51^, may remove spurious findings. To do this, we simulate null relationships between heteroplasmy levels and a donor phenotype, donor age, using realistic ranges of RNAseq coverage, VAFs at apparent mtDNA heteroplasmy, and donor age relationships with RNAseq coverage we obtain from the GTEx data (**Supplementary Methods, Extended Data Figure 2, Supplementary Table S7**). We then ask whether the LM or the BB model are well calibrated in assessing the simulated relationship between VAFs and donor age at different simulated RNAseq coverages, through comparing their false positive (FP) rates (**Supplementary Methods**). We find that while the BB model is well calibrated across the whole realistic range of parameters we assess (**Extended Data Figure 2**), LM tests becomes inflated (FP > 0.05) at low coverages (coverage = 50), especially when combined with a high level of heteroplasmy and a decreasing relationship between coverage and donor age. Where both models show inflation, BB always performs better than LM. As such, we derive a two-step heteroplasmy association testing procedure that first identifies putative significant associations using a LM, then verifies them using a BB model, thereby combining both the runtime advantage of LMs and the rigour of BB with permutations (**Methods, Extended Data Figure 2**). We apply this testing procedure for all apparent mtDNA heteroplasmy association testing in this paper.

### Effect of donor age on mtDNA heteroplasmy and mtRNA modifications

Using the two-step heteroplasmy association testing procedure, we first investigate the tissue-specific relationship between donor age and apparent mtDNA heteroplasmy at 556 apparent heteroplasmic sites (26-319 per tissue, **Methods**). We find 109 significant associations with donor age at study-wide significance (**Figure 3A, Supplementary Table S8**), the majority of which (95.4%) are at mtDNA heteroplasmy. All of the donor age effects on mtDNA heteroplasmy and mtRNA modifications are positive, consistent with previous findings of age-related increase in mtDNA heteroplasmy levels in blood^30,31^.

**Figure 3:**
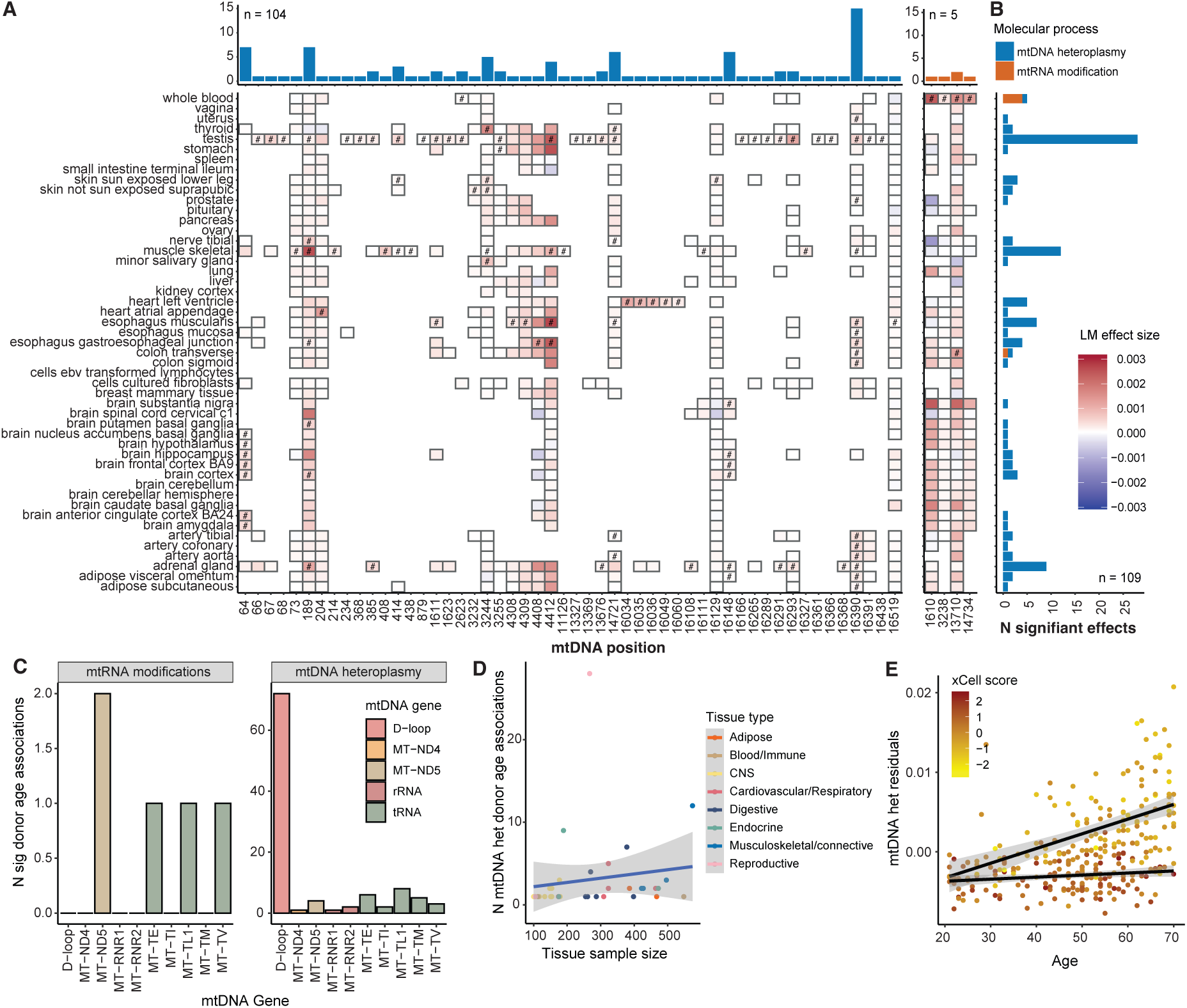
**A.** Top: number of tissues in which mtDNA heteroplasmy (blue, total n=104) and mtRNA modifications (orange, total n = 5) are associated with donor age per site; bottom: heatmap of effect sizes from a linear model testing for associations between donor age and mtDNA heteroplamsyt and mtRNA modifications per position per tissue; boxes are coloured by effect sizes; significant effects are marked with #. **B.** Barplot of number of apparent heteroplasmy per tissue associated with donor age; bars are coloured by molecular process: mtDNA heteroplasmy (blue) or mtRNA modification (orange). **C.** Barplots showing numbers of significant donor age associations identified in mtRNA modifications and mtDNA heteroplasmy across tissues per mtDNA gene; bars are coloured by mtDNA gene. D. Scatterplot showing the number of significant mtDNA heteroplasmy associations with donor age plotted against the sample size in the tissues they are identified in; each dot represents a tissue; dots are coloured by tissue; blue line indicates linear relationship between number of significant associations and tissue sample size. **E.** Scatter plot showing the only significant and negative interaction effect between donor age and xCell enrichment score of Epithelial Cells in Colon Transverse at mt.16390 (P value = 2.11 x 10^-8^); we show donor age on the x axis, the residuals of mtDNA heteroplasmy VAF corrected for sex and gPCs on the y axis, and the xCell scores of the tested cell type in colour; the black two lines show the linear relationships between donor age and VAF at mt.16390 at high (>mean) and low (<=mean) xCell scores, for visualization of the difference in effects at high and low xCell scores.

We find that the number of donor age associations with VAFs at apparent mtDNA heteroplasmy per tissue is not correlated with tissue sample size (Pearson r = 0.14, P = 0.43) or median tissue mtDNA-CN^48^ (Pearson r = 0.03, P = 0.85). Rather, we find the greatest number of donor age effects on mtDNA heteroplasmy in specific tissues including testis (n = 28), muscle skeletal (n = 12), and adrenal gland (n = 9, **Figure 3B**). The strongest effect across all tissues is seen at mt.4412 in Esophagus Muscularis where there is an increase in VAF by 2.50% for every 10 years increase in age. We find that donor age effects on mtDNA heteroplasmy at 15 positions are shared across multiple tissues: most prominently, mtDNA heteroplasmy at mt.16390 and mt.189 are significantly associated with donor age in 15 and 7 tissues from different organ systems, while mtDNA heteroplasmy at mt.64 is significantly associated with donor age in 7 CNS tissues (**Figure 3A**). Finally, we obtain replication of the donor age association signal in peripheral blood mononuclear cells (PBMCs) at mt.2623 using mitochondrial single-cell assay for transposase accessible chromatin by sequencing (mtscATACseq)^19,52,53^ (mt.2623A>G Pearson r=0.788, **Extended Data Figure 3, Supplementary Methods**). Interestingly, though most of the donor age associations are with VAFs at mtDNA heteroplasmy, most of them are on mtDNA encoded tRNAs and non-coding regions, rather than on protein-coding genes (**Figure 3C**). For mtRNA modifications, we find the greatest number of significant donor effects in Whole Blood (n = 5, **Figure 3B**).

Overall, we do not observe significant associations between the number of donor age associations with mtDNA heteroplasmy and tissue sample size (too few of mtRNA modification associations for testing across tissues), suggesting differences between tissue findings are not merely due to power, but potentially tissue-specific mechanisms, especially as seen in the high number of associations in the Testis (**Figure 3D**). For mtRNA modifications, we observe a consistent pattern of donor age associations with large effect sizes in CNS tissues that is absent in other tissue types, none of which are individually significantly associated with donor age (**Figure 3A**). Nonetheless, the consistent donor age effects on mtRNA modifications across CNS tissues is unlikely to have arisen by chance (sign test P = 2.84×10^-14^), and we speculate they have potential consequences on CNS mtDNA gene regulation.

We further asked if the donor age associations with apparent mtDNA heteroplasmy are cell-type specific within tissues, through testing for interaction effects between xCell enrichment scores of 45 highly enriched cell types in 39 GTEx tissues^54^ and donor age on VAFs at apparent mtDNA heteroplasmic sites (**Methods**). We find only one significant and negative interaction effect between donor age and xCell enrichment score of Epithelial Cells in Colon Transverse at mt.16390 (P value = 2.11 x 10^-8^, **Figure 3E**), suggesting the donor-age association at this position in Colon Transverse is driven by cell types other than Epithelial Cells.

### Effect of apparent mtDNA heteroplasmy levels on tissue-specific mtDNA gene expression

We then investigate if apparent mtDNA heteroplasmy VAFs are associated with mtDNA gene expression levels. We use the same two-step association framework to test for associations between mtDNA gene expression and both VAFs at mtDNA heteroplasmy and mtRNA modification sites, this time modeling donor age as an additional covariate (**Methods**). We find 784 significant associations at study-wide significance across 48 tissues (**Methods**), of which 576 are mtDNA heteroplasmy and 208 are mtRNA modifications (**Figure 4A, Supplementary Table S9**). The number of significant associations vary greatly between tissues, with the highest number of effects identified in Cell Cultured Fibroblasts (193) followed by Whole Blood (95), with Brain Amygdala (1) and Artery Coronary (1) showing the fewest effects (**Figure 4B**). This is likely partly due to power differences for association testing between tissues: the number of significant effects found for mtDNA heteroplasmy (Pearson r = 0.50, P = 9.70×10^-4^) and mtRNA modifications (Pearson r = 0.42, P = 6.91×10^-3^) are both significantly correlated with sample sizes per tissue (**Figure 4C**).

**Figure 4:**
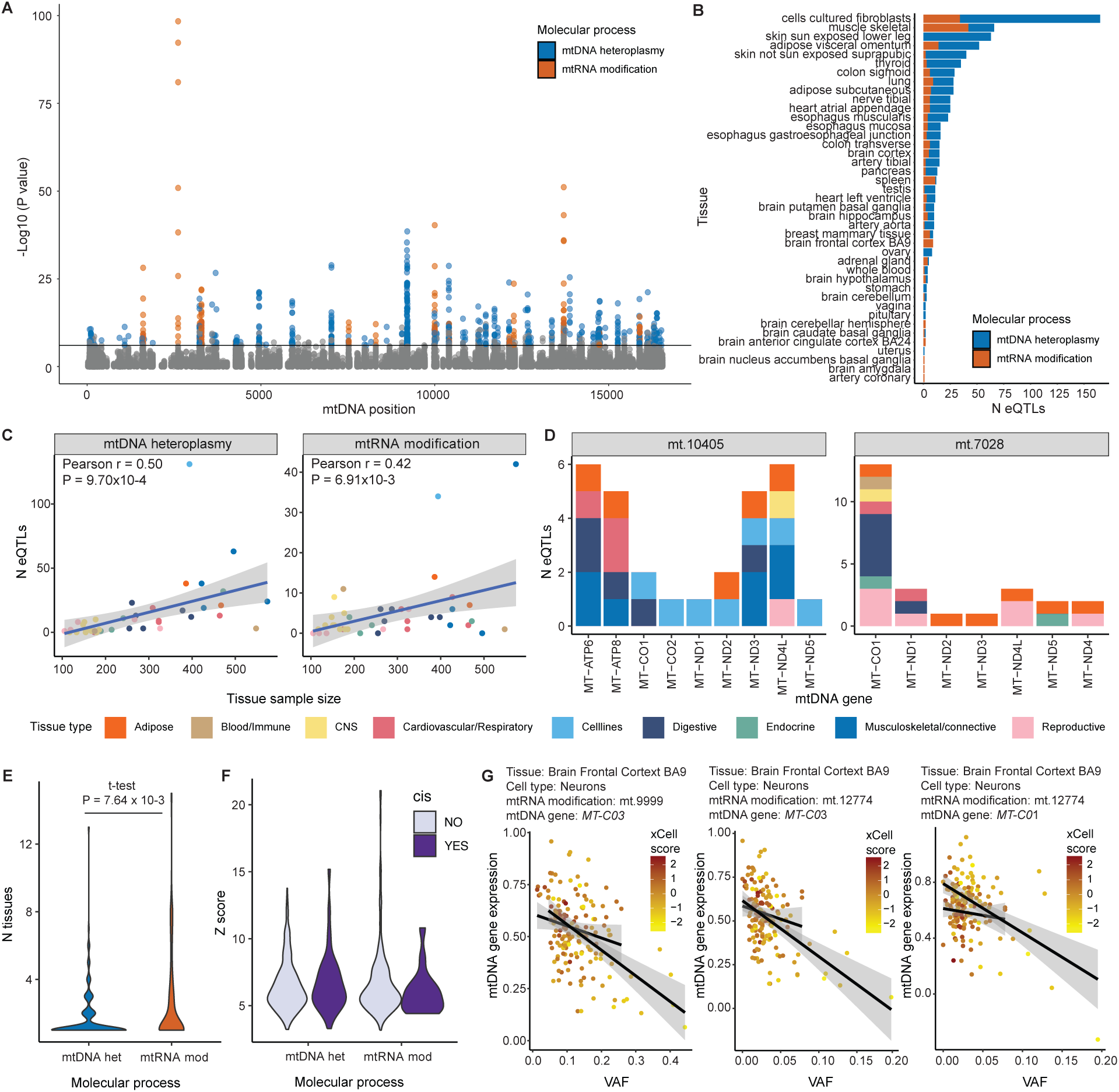
**A.** Manhattan plot showing the significance of all apparent heteroplasmy eQTLs associations with mtDNA genes across tissues; each dot represents an apparent heteroplasmic site; dots are coloured by the molecular process; study-wide significance threshold for the linear model (LM) test is shown in a black line. **B:** Barplot showing the number of significant apparent heteroplasmy eQTLs per tissue; bars are coloured by molecular process. C. Scatterplot showing the number of mtDNA heteroplasmy and mtRNA modification eQTLs per tissue, plotted against tissue sample size; each dot represents a tissue; dots are coloured by tissue type. **D.** Barplots showing the numbers of eQTLs identified at mt.10405 (MT-TR) and mt.7082 (MT-CO1) per mtDNA genes in different tissues; bars are coloured by tissue categories. **E.** Violin plots showing the number of tissues with the same eQTLs identified in mtDNA heteroplasmy and mtRNA modifications per gene, and a t-test showing there are more shared mtRNA modification effects between tissues on mtDNA gene expressions across tissues than mtDNA heteroplasmy effects (t-test P = 7.64 x 10^-3^); violins are coloured by molecular process. **F.** Violin plots showing the Z scores of eQTL effects at cis- and non-cis mtDNA genes identified at mtDNA heteroplasmy and mtRNA modification sites; violins are coloured by whether the mtDNA gene is in cis with the variation tested. **G.** Three examples of significant interaction effects on tissue-specific mtDNA gene expression levels between VAFs at apparent mtDNA heteroplasmy and xCell scores of enriched cell types in Brain Frontal Cortex BA9^34^; for each significant interaction we show the VAF of the apparent mtDNA heteroplasmy on the x axis, the residuals of log(TPM+1) values for gene expression levels of mtDNA genes (corrected for PEER factors calculated per tissue) on the y axis, the xCell scores of the tested cell type in colour, and the black two lines show the linear relationships between the VAF and mtDNA gene expression levels at high (>mean) and low (<=mean) xCell scores, for visualization of the difference in effects at high and low xCell scores.

Most of the significant associations are found between mtDNA gene expression levels and mtDNA heteroplasmy (73.47%) rather than mtRNA modifications. In particular, mtDNA heteroplasmy at mt.10405T>A/G (in *MT-TR*) and mt.7028T/C>C/T (in *MT-CO1*) show the greatest numbers of associations, being associated with the expression levels of 9 and 7 genes respectively (**Figure 4D**). Interestingly, while the effects of mt.7028 are consistently negative across almost all genes it has a significant association with, those for mt.10405 are varied (**Extended Data Figure 4**), raising the question if mtDNA effects in mt-tRNAs are more likely to have gene-specific regulatory effects. Overall, we find more shared mtRNA modification effects on mtDNA gene expressions across tissues than mtDNA heteroplasmy effects (t-test P = 7.64 x 10^-3^, **Figure 4E**). Finally, 7.12% of mtDNA heteroplasmy eQTLs are in genes where the mtDNA heteroplasmy is located, and 7.69% of mtRNA modification eQTLs are in genes right next to genes the mtRNA genes the modifications are found in. We call these “cis” eQTLs, and ask if the effects at these “cis” eQTLs are stronger than those from “non-cis” eQTLs. We do not find any significant difference in z-scores between “cis” and “non-cis” eQTLs for either (**Figure 4F**).

Like we did for donor-age associations, we ask if gene expression associations with VAFs at apparent mtDNA heteroplasmy are cell type-specific, through testing for interaction effects between xCell scores of enriched cell types in each tissue with VAFs at apparent mtDNA heteroplasmy (**Methods**). We find a total of 18 significant interactions, including 15 at mtDNA heteroplasmy and 3 at 2 mtRNA modification sites (**Figure 4G**, **Extended Data Figure 5**, **Supplementary Table S10**).

### Shared effects on donor age and MT gene expression mediated by heteroplasmy levels

We find that the VAFs at 10 tissue-specific apparent mtDNA heteroplasmy sites are significantly associated with both donor age and expression of mtDNA genes in the same tissue (**Supplementary Table S11**). All 10 apparent heteroplasmy sites are in the D-Loop or in mt-tRNAs, though 9 are mtDNA heteroplasmy, and only 1 is a mtRNA modification site at mt.14734 in Whole Blood (*MT-TE* mt-tRNA for glutamic acid, transcript position 9).

For all 10 sites, we ask if the associations between apparent mtDNA heteroplasmy levels and mtDNA gene expression are in fact mediating donor age effects on the latter. To do this, we perform a mediation analysis using partial correlation of residuals among apparent mtDNA heteroplasmy levels, donor age, and mtDNA gene expression levels^32,33^ (**Figure 5A**, **Supplementary Methods**). We find that across 10 sites, 9 of them support our hypothesis that apparent mtDNA heteroplasmy mediated the effect of donor age on mtDNA gene expression levels, including the mtRNA modification at mt.14734 (**Figure 5B**, **Supplementary Table S11,S12**). Notably, the relationship between donor age and apparent mtDNA heteroplasmy VAFs, conditional on mtDNA gene expression levels, is always positive, consistent with our expectation of donor-age related accumulation of mtDNA heteroplasmy in 9 out of the 10 sites. In contrast, the relationship between apparent mtDNA heteroplasmy VAFs and mtDNA gene expression levels, conditional on donor age, are sometimes negative, suggesting some mtDNA heteroplasmy levels result in reduced gene expression levels.

**Figure 5:**
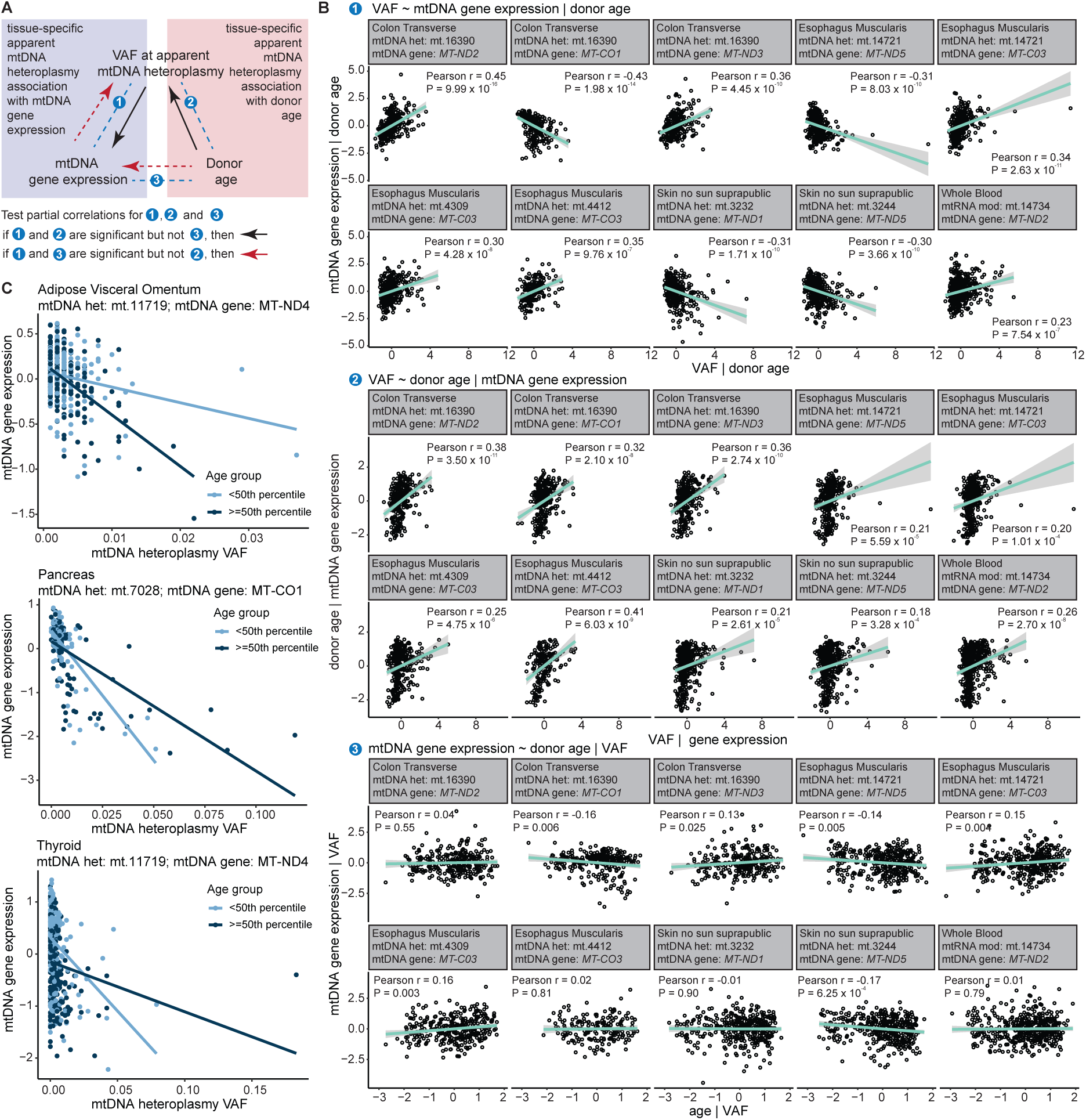
**A.** Schematic showing the three partial relationships we test between VAF, donor age and mtDNA gene expression in order to identify potential causal relationships between them; arrows show hypothesized direction of causal effects. **B.** Partial correlations at 10 apparent heteroplasmy sites where there is both a significant eQTL association and donor age association; top: partial correlations between VAF and mtDNA gene expression regressing out donor age from both; middle: partial correlations between VAF and donor age regressing out mtDNA gene expression from both; bottom: partial correlations between mtDNA gene expression and donor age regressing out VAF from both; dot represents values in a sample; green line represents the Pearson correlation, with both r and P values indicated in the plots. **C.** Three examples of significant interaction effects on tissue-specific mtDNA gene expression levels between VAFs at apparent mtDNA heteroplasmy and donor age. For each significant interaction we show the VAF of the apparent mtDNA heteroplasmy on the x axis, the residuals of log(TPM+1) values for gene expression levels of mtDNA genes (corrected for PEER factors calculated per tissue) on the y axis, the donor age groups (<50TH percentile and >= 50 percentile) in colour, along with lines of the respective colours showing the linear relationships between the VAF and mtDNA gene expression levels at high (>=50 percentile) and low (<50th percentile) donor ages, for visualization of the difference in effects at high and low donor ages.

In addition, we ask a separate question of whether the associations between mtDNA heteroplasmy effects and mtDNA gene expression are different at different ages. To this end, we test for interaction effects between apparent mtDNA heteroplasmy VAFs and donor age at the 784 significant associations between apparent mtDNA heteroplasmy VAFs and mtDNA gene expression levels (**Methods**). We find 11 significant interaction effects between donor age and apparent mtDNA heteroplasmy VAFs on apparent mtDNA gene expression levels, all of which are at mtDNA heteroplasmy sites, not mtRNA modifications. None of them overlap the 9 instances where mtDNA heteroplasmy mediated the effect of donor age on mtDNA gene expression levels (**Figure 5C, Extended Data Figure 6**, **Supplementary Table S13**).

### Effect of mtRNA modifications on transcript processing

The m1A/G methylation of mt-tRNAs at position 9 (p9) has been found to be carried out by a protein complex TRMT10C–SDR5C that not only acts as a methyltransferase, but also as a pre-mt-tRNA chaperone that sequentially recruits and assembles the endonuclease complex that cuts the mt-tRNA from the mRNA transcript on its 5’ end, essential for the maturation of the 5’ protein-encoding mRNA and further pre-mt-tRNA processing at the 3’ end^26,55–58^ (**Figure 6A**). As such, one study has previously tested the hypothesis that p9 mt-tRNA modification levels regulate mRNA levels 5’ of the tRNA^22^, using RNAseq data on blood cell types, including Whole Blood in GTEx v6^59^.

**Figure 6:**
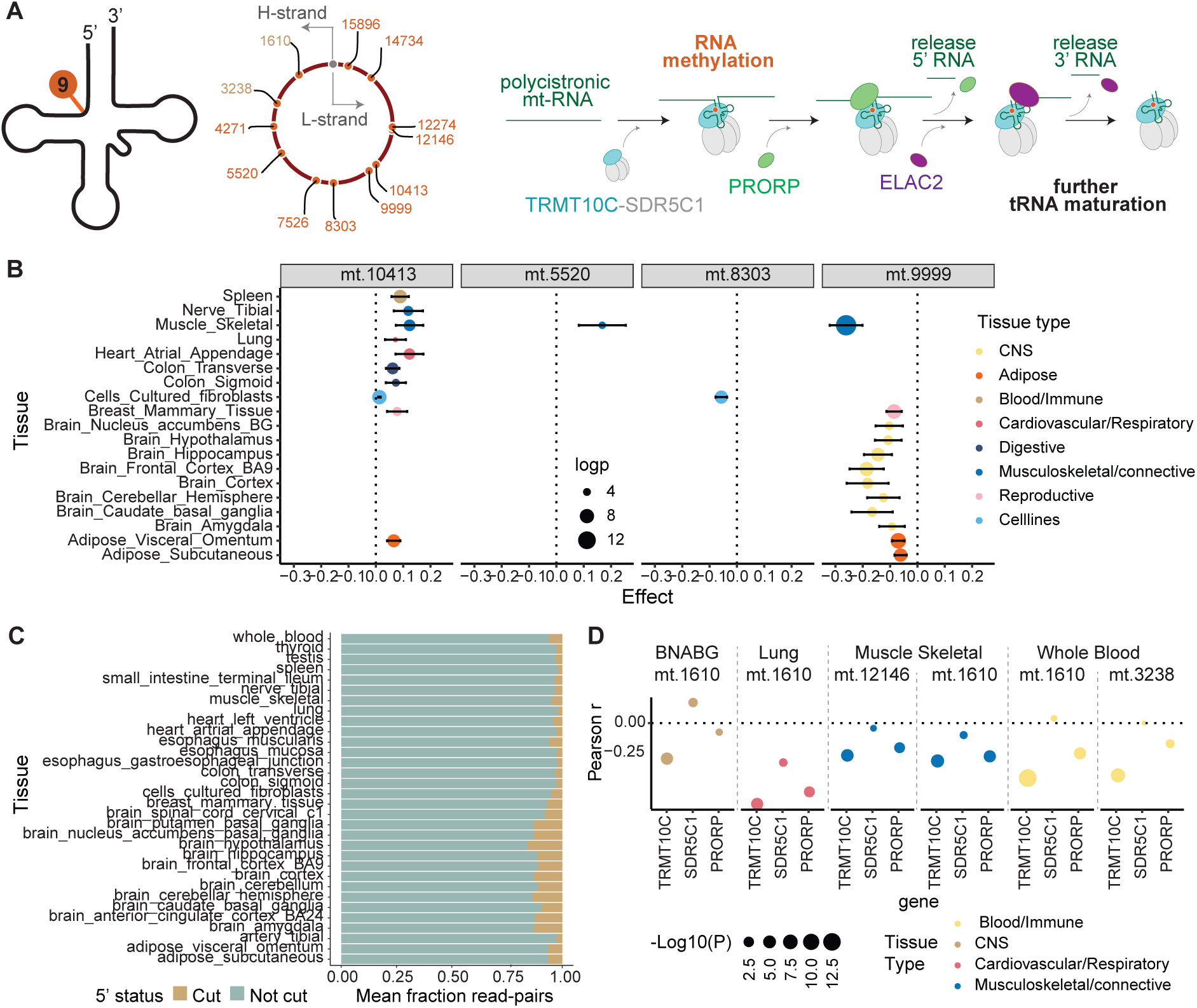
**A.** Left: Schematic showing the p9 methylation on mt-tRNAs identified in all samples and tissues in GTEx on both the H-strand (yellow) and the L-strand (orange); right: a schematic showing a simplified methylation process at these p9 methylation sites catalysed by a protein complex made of TRMT10C and chaperone SDR5C1, which then recruits endonuclease PRORP for releasing 5’ RNA from the methylated mt-tRNA, leading to further mt-tRNA maturation and release at the 3’ end by ELAC2. **B.** Plot showing the effect of VAF at four p9 methylation sites previously reported to be associated with gene expression levels of their respective 5’ mtDNA genes, now tested in GTEx v8 tissues; each dot represents a tissue; sizes of dots are representative of the -log10(P) of the associations; error bars indicate the 95% CI of the effects; dots are coloured by tissue type; we replicate previous findings and find more associations for these 5’ mtDNA genes. **C**. Bar plots showing the mean fraction of read-pairs with p9 modifications in each tissue identified directly from QC-ed RNAseq data that are likely cut or not cut at the 5’ ends; bars are coloured by their putative cut status. **D.** Pearson r between VAF at p9 modifications and log(TPM+1) gene expression levels of components of the mt-tRNA processing pathway in tissues where there is at least one significant correlation, at p9 positions mt.1610 and mt.12146 where there is at least one significant correlation across all tissues; each dot represents a tissue, dots are coloured by tissue and sizes of dots represent the -log10(P) of the correlation.

Here we ask if we can replicate the findings from this previous study and derive more biological insights using VAF at mt-tRNA modifications sites and more detailed modification features derived from raw RNAseq read data from all tissues in GTEx v8 (**Methods**, **Supplementary Methods**). We first perform the same association analysis between VAF of mtRNA modifications with 5’ gene expression at the 7 p9 mt-tRNA modification sites with a protein coding 5’ gene as the previous study did^22^, using data in GTEx v8 Whole Blood. Our results replicate the only previously found significant association between p9 *MT-TG* modification at mt.9999 and the expression of its 5’ gene *MT-CO3* in GTEx v6 and CARTaGENE, with the same direction of effect (**Supplementary Table S14**). We then perform the same analysis in all other tissues in GTEx v8, and identify 24 significant associations between the VAF at p9 mt-tRNA modifications and their 5’ gene expressions in 19 tissues at study-wide significance (VAF-based analysis, **Figure 6B**, **Supplementary Table S15**). We find that effects of different p9 mt-tRNA modifications on 5’ gene expressions are variable, suggesting the expected coupling of the methylation and the 5’ “cutting” process previously reported^22,25^ may not be that straightforward.

We then take the analysis one step further, asking if the p9 mt-tRNA modifications are truly affecting 5’ gene expression levels through the simultaneous recruitment of endonucleases for a mtRNA “cut” at the 5’ end of the mt-tRNA. We therefore quantify the number of read pairs supporting both a p9 mt-tRNA modification effect and a 5’ “cut”, as well as those only supporting p9 mt-tRNA modification in all samples per tissue (**Figure 6C**, **Supplementary Figure S5**, **Methods**). We then ask if gene expression levels of the 5’ gene are associated with a 5’ “cut” among p9 modified reads. We find a total of 21 significant associations between the 5’ “cut” and the 5’ gene expression levels in this read-based analysis study-wide. 19 out of the 24 significant VAF-based associations with 5’ gene expressions are significant in this read-based analysis (**Supplementary Table S15**), meaning a majority of VAF effects of mt-tRNA modifications on the 5’ gene expression may indeed be attributable to a “cut” on the 5’ end of the mt-tRNA.

Across all tissues, effects of the same p9 mt-tRNA modifications on 5’ gene expression are all in the same direction. However, we find that the directions of effect can be different between p9 mt-tRNA modifications, even though they are all, in theory, methylated by TRMT10C^23–26^. We find that the VAFs of p9 mt-tRNA modifications are only significantly correlated with the inverse normalized log(TPM +1) gene expression levels of *TRMT10C* in a few tissues (**Methods**), and in those significant cases, the correlations are invariably negative (**Figure 6D**). This is counter-intuitive given TRMT10C^23–26^ is previously established to both mediate these methylations and the recruitment of the endonuclease complex (**Figure 6A**). Furthermore, we find that the VAFs of these p9 mt-tRNA modifications, found to be significantly correlated with *TRMT10C* gene expression in the same tissues, have variable relationships with the gene expression levels of the chaperone gene *SDR5C1* or the catalytic ribonuclease *PRORP* in different tissues (**Figure 6D**). This is consistent with our findings of both positive and negative associations between p9 mt-tRNA modifications and 5’ gene expression levels (**Figure 6B**, **Supplementary Table S15**). We think these between-site and between-tissue differences may be due to complex, feedback-based, and potentially redundant regulatory processes that compel further investigations into all components of this machinery in a tissue-specific manner.

## Discussion

In this study, we comprehensively characterize the landscape of common mtDNA heteroplasmy and mtRNA modifications across 49 human tissues assayed with bulk RNAseq in GTEx v8. As mtDNA heteroplasmy and mtRNA modifications are tissue-specific, we aim to establish a robust framework for identifying and testing both types of variations from tissue-specific RNAseq data.

First, we find strand bias highly commonplace in RNAseq data on the mtDNA transcripts (**Supplementary Figure 4**), especially those for mt-tRNAs and mt-rRNAs which are likely off-target reads in polyA+ selected libraries in GTEx^32,60^. This necessitates modifications to previously proposed pipelines optimized for filtering out strand biases in mtDNA variant identification from WGS data^33,61^. Second, we find that VAFs at apparent mtDNA heteroplasmic sites show significant heteroskedasticity across RNAseq coverages, which means associations between heteroplasmy and donor phenotypes that affect RNAseq coverage (age, gene expression levels) are confounded. We therefore adapt the beta-binomial model commonly used for ASE^40,51^ testing to account for this heteroskedasticity, enabling further investigations into relationships between apparent mtDNA heteroplasmy and donor phenotypes. We see both these improvements as important contributions to the development of a protocol for mtDNA heteroplasmy investigations. As our donor age association in Whole Blood is replicated using cell-type specific mtDNA heteroplasmy data obtained from mtscATACseq^19,52,53^, we deem our approaches to be of considerable relevance as the community shifts from using bulk RNAseq to single-cell data.

Employing the new association test framework, we present the first comprehensive investigation into the relationship between donor age and tissue-specific mtDNA heteroplasmy and mtRNA modification levels. We observe the strongest effect at mt.4412 in Esophagus Mucosa, where there is an increase of ∼15% in mtDNA heteroplasmy over 50 years. We identify many significant associations with the former, consistent with established understanding that mtDNA mutations accumulate with age^16^, but not many in the latter, which may be due to both low power and a more stringent control of mtRNA modification levels due to their importance in the mtRNA transcript processing pathways. Though there are many proposed mechanisms for how the levels of mtDNA heteroplasmy or mtRNA modification are moderated to ensure they do not exert detrimental effects on mtRNA transcripts (where mtDNA heteroplasmy are transcribed) or on gene expression regulation (where mtDNA heteroplasmy or mtRNA modifications play regulatory roles), the exact mechanism, likely context dependent, remains to be determined. One way forward would be to overlay our findings of donor age relationships with mtDNA heteroplasmy or mtRNA modifications with tissue-specific genetic regulation of key components of the regulatory pathways with previous evidence of keeping both variations in check.

Similarly, we present the first comprehensive and tissue-specific assessment of mtDNA heteroplasmy and mtRNA modification effects on mtDNA gene expression. For mtDNA heteroplasmy, we find that a majority of the variations that show both significant donor age and gene expression associations (in the same tissue) are likely to mediate the donor age effects on gene expression, potentially accounting for the effect of aging on mitochondrial function. Overall, we find much fewer mtDNA heteroplasmy effects on mtDNA gene expression levels than homoplasmic effects (Gianoullis et al. *related submission*). This may be partly because we have much lower power to detect the smaller effects heteroplasmic variations are likely to exert on gene expression levels. However, it is also possible that compensation mechanisms result in a much lower level of penetrance of heteroplasmic effects on organismal phenotypes than homoplasmic variations^62^. One interesting direction for future investigations, given mtDNA heteroplasmy mediate age effects, is whether these compensatory mechanisms and the buffers they create become thinner as one ages. For example, it is established that in general mtDNA-CN becomes lower with aging, which may result in its gene expressions becoming more stochastic^50,63,64^.

For mtRNA modifications, we make an attempt to assess the direct effect of p9 mt-tRNA modifications on the gene expression levels of their 5’ genes, given previous findings that the protein TRMT10C involved in methylation of these sites also have roles in the recruitment of the 5’ transcript processing ribonuclease. Though our findings replicate and extend previous findings of significant relationships between p9 mt-tRNA modifications and 5’ gene expression, further analyses involving the tissue-specific gene expression levels of the methylase *TRMT10C*, the ribonuclease *PRORP* and the chaperon *SDR5C1* have revealed a more complex and tissue-specific picture of regulatory mechanisms that have hitherto not been taken into account. Future analyses into these mechanisms may therefore involve modeling of the gene expression coordination between these regulatory components, and their effects on both mt-tRNA modifications and 5’ gene expression levels, as well as the inclusion of data generated by more targeted assays^65,66^. Further, mtRNA modifications may affect mitochondrial protein translation in addition to transcript processing^67^, and there is early work on quantifying mt-tRNA modifications with Nanopore sequencing^65,66^. These hold promise for the investigation on the functional consequences of the full repertoire of mt-tRNA modifications on mitochondrial protein levels and downstream physiology^67^.

Though we do not touch upon the implication of our findings in health and disease, our methodologies and findings may provide a basis for examining disease associations with mtDNA heteroplasmy and mtRNA modifications^68,69^, through providing a tissue-specific reference atlas derived from a general (rather than a disease) population. Similarly, previously identified disease associations with the machinery of mt-tRNA modifications^70^ can be, we propose, modeled together with mt-tRNA modifications effects on gene expression levels going forward. In any case, establishing direct causal connections between mtDNA heteroplasmy or mtRNA modification associations with diseases remains very difficult, since most associations are performed in separate datasets, and currently there are no methods established for inferring causality of mtDNA variants like there is for nucDNA variants (e.g. Mendelian Randomization^71^). Actual causal inference may have to be made through assessing mtDNA variants in individuals with diseases^72^, or through engineered mtDNA base-editing in cellular or animal models^73^.

Finally, our investigations in this paper are limited to individuals of European genetic ancestry. While we expect mtDNA homoplasmic variants to differ in effect widely across populations, due to their vastly different allele frequencies across populations and their mtDNA Haplogroups, whether and how mtDNA heteroplasmy or mtRNA modification differ in their position or effect in between populations remains to be investigated. We suspect these effects may in fact be more universal than those of homoplasmic variations. Increasing efforts to collect data across diverse ancestries will enable this research in the future.

Overall, our work characterises mtDNA heteroplasmy and mtRNA modifications in human tissues, advances our understanding of their relationships with age and mtDNA gene expression levels, and forms the basis for further investigations into their phenotypic relevance.

## Methods

### GTEx mtDNA heteroplasmic variant calling

For identifying mtDNA variants in the 684 GTEx v8 samples with European descent (**Supplementary Methods**), we extracted reads mapping to the rCRS mitochondrial reference genome (NC_012920) from 12,577 RNA sequencing samples (RNAseq) 49 primary tissues and 2 cell lines from 684 donors of European descent, as well as whole-genome sequencing samples (WGS) obtained from Whole Blood of the same 684 donors. We filtered the mtDNA reads from both RNAseq and WGS on read alignment flags (removing unmapped reads, unmapped pairs, PCR duplicates, secondary mapping and reads failing vendor QC; using the -F 3852 option) with samtools^74^ (v1.9). We then called both homoplasmic (where there is only 1 allele at a position supported by sequencing reads) and heteroplasmic (where there are more than 2 alleles per position supported by sequencing reads) mtDNA variants from the filtered reads using mtdna-server^23^ (using options type==2, status == PASS). We then apply a set of additional filters to account for strand bias in RNAseq derived heteroplasmy calls (**Supplementary Methods, Supplementary Table S1**). We provide a summary of all apparent mtDNA heteroplasmic variants obtained from RNAseq in **Supplementary Table S2**. For all heteroplasmic variations we identify from WGS and RNAseq, we assume the major allele identified at the same position in WGS in the same donor is the inherited allele, while the other alleles are due to somatic mutations in the mtDNA or modifications in the mtRNA. We call the latter heteroplasmic allele, and its frequency in the sample variant allele frequency (VAF).

### Selection of mtDNA heteroplasmic sites for association testing

For each apparent mtDNA heteroplasmic position we use VAFs as genotypes of samples with identified heteroplasmy at the position in individuals with greater than 10 reads on both the forward and reverse strands. We set genotypes of those samples with fewer reads as missing (NA) and only keep samples with > 75 % non-missing values. We only perform association testing in tissues where there are more than 60 samples and in which > 75 % of the samples have non-missing values for a particular mtDNA heteroplasmy position, and at positions where there is non-zero variance in heteroplasmy among samples.

### Phenotype association testing

In our two-step testing procedure for the association between heteroplasmic levels and donor phenotypes, we first test each apparent heteroplasmy passing quality filters using a linear model adjusted for sex and the first 5 genetic principal components (gPC, **Supplementary Methods**) as covariates: V = β_0_ + β_1_F + β_2_A + e, where V is VAF of the apparent heteroplasmy, F the covariates, A the donor phenotype, and e error term. For all tests that pass the study-wide Bonferroni-corrected P-value threshold of 0.05, we then perform the same association test again using the Beta-Binomial (BB) model, now modelling V as (N_Vreads_, N_Oreads_), where N_Vreads_ is the number of reads supporting the variant alleles, and N_Oreads_ is the number of reads supporting the other allele. For this we use the betabin() function from the aod R package v.1.3.3. Further, we perform 10,000 permutations of the BB model for each apparent heteroplasmy passing study wide BB Bonferroni correction (< 0.05) and obtain the empirical P-values as (N_permutation_pvals_ < N_analytical_pvals_)/N_permutations_. We then perform a study-wide Bonferroni correction on the empirical P-values to obtain the final significant associations.

### Apparent heteroplasmy cis-eQTLs on mtDNA encoded genes

In testing for apparent heteroplasmy effects on mtDNA encoded genes, we use residuals of log(TPM+1) values for gene expression levels of mtDNA genes, corrected for PEER factors calculated per tissue (**Supplementary Table S2**), as donor phenotype in the above two-step testing scheme. We also include donor age, in addition to sex and gPCs, as a covariate. Notably, we test for apparent heteroplasmy cis-eQTLs only in 12 mtDNA encoded, protein-coding and polyadenylated genes^75^, unlike in other work that test for mtDNA cis-eQTLs and nuclear DNA trans-eQTLs in all 13 mtDNA-encoded protein coding genes^34^. Further, though we characterise apparent heteroplasmy in 49 tissues, we only test 48 out of 49 tissues for associations between VAFs of apparent mtDNA heteroplasmy and mtDNA gene expression levels, not Kidney Cortex (N=64), consistent with mtDNA eQTL analyses on GTEx v8^34^ on and previous recommendations^76^.

### Deconvolution of associations signal by cell type

We perform cell-type interaction analysis to identify donor age effects on apparent heteroplasmy VAF and gene expression association driven by specific cell types (adipocytes, epithelial cells, hepatocytes, keratinocytes, myocytes, neurons, neutrophils). We filter for cell types with median GTEx-provided xCell enrichment scores^54^ > 0.1 for each tissue, as shown in a recent study on mtDNA gene regulation using GTEx v8 data^34^. eQTLs identified in tissues where no cell types pass the filter are not tested for cell-type interaction effects^54^. For investigating donor age effects on apparent heteroplasmy VAFs, we obtain interaction effect between donor age and xCell scores of cell types passing the filter per tissue using the model introduced in Westra et al.^77^: V = β_0_ + β_1_F + β_2_A + β_3_C_celltype_ + β_4_A⋅C_celltype_ + e, where F consists of covariates including sex and 5 gPCs, V is VAF of the apparent heteroplasmy, A the donor age, C the cell-type proportion, CxA the interaction term between cell-type score and donor age, and e the error term. For investigating apparent heteroplasmy VAF effects on gene expression levels, we use: GE = β_0_ + β_1_F + β_2_V +β_3_C_celltype_ + β_4_V⋅C_celltype_ + e, where GE is the residuals of mtDNA gene expression levels log(TPM+1) after controlling for tissue-specific numbers of PEER factors^34^. For each analysis, we only test interaction effects at significant associations between donor age and apparent heteroplasmy VAF or apparent heteroplasmy VAF and gene expression levels respectively. To control for multiple testing, we perform Bonferroni correction across all interaction effects tested in each analysis at study-wide levels. We consider all interaction effects with adjusted P < 0.05 at study-wide level to be significant.

### Interactions between VAFs at apparent mtDNA heteroplasmy and donor age

For investigating the interaction effects between apparent heteroplasmy VAF and donor age on mtDNA gene expression levels, we use: GE = β_0_ + β_1_F + β_2_V + β_3_A + β_4_V⋅A + e, where GE is the residuals of mtDNA gene expression levels log(TPM+1) after controlling for tissue-specific numbers of PEER factors, F consists of covariates including sex and 5 gPCs, V is VAF of the apparent heteroplasmy, A the donor age, VxA the interaction term between VAF of the apparent heteroplasmy and donor age, and e the error term. We perform Bonferroni correction across all interaction effects tested at study-wide level and consider all interaction effects with adjusted P < 0.05 to be significant.

### Inferring mitochondrial transcript processing status

To extract reads at all 12 p9 mt-tRNA modification sites identified in all samples across tissues, we use pysam^74,78^ (v0.22.0) pileup, retaining only paired reads with mates mapped to the mtDNA reference with mapping quality score > 20, and excluding duplicate reads, as well as secondary and supplementary alignments. For the aligned bases at each p9 mt-tRNA modification, we require base quality score > 20 and alignment quality score > 30. We only retain samples with more than 10 read-pairs with aligned base at p9 mt-tRNA modification site passing quality control for analysis. Here we further exclude samples where the Spearman correlation between the proportion of methylated read-pairs and their previously estimated VAF is below 0.75 (**Supplementary Figure S5**). For each read-pair with aligned base at p9 mt-tRNA modification site passing quality control, we determine its mt-tRNA 5′ and 3′ processing status by evaluating whether the start coordinate of its first read was smaller than the 5′ end or larger than the 3′ end of each mt-tRNA gene, accounting for H- and L-strandedness of the respective mtDNA genes.

### Mitochondrial transcript processing analysis

We first test for the relationship of mtRNA methylation levels and 5’ protein coding gene expression at those p9 mt-tRNA modification sites where the 5’ gene was immediately upstream, as previously described^22^, using linear regression: V = β_0_ + β_1_F + β_2_GE + e, where V represents the VAF at p9 mt-tRNA modifications, F consists of covariates including donor sex, age, and the first five genomic principal components (gPC1–5), GE is the residuals of the 5’ gene expression log(TPM+1) adjusted for tissue specific PEER factors, and e represents the error term. Next, we assess whether the number of methylated and putatively cut read-pairs is associated with 5’ protein coding gene expression using a binomial regression, specified as (N_mod,cut_, N_mod,not-cut_) = β_0_ + β_1_F + β_2_GE + e. We specified a null model excluding β_2_GE and performed a likelihood ratio test to obtain p-values using the lmtest R package (v.0.9-40). For each step, we use a study-wide Bonferroni threshold (adjusted P < 0.05) for multiple-testing correction.

## Acknowledgements

SW is supported by the Joachim Herz Foundation. SW and XG are supported by the Munich School for Data Science (MUDS). PaK and LSL are supported by the Hector Fellow Academy. LSL is supported by an Emmy Noether fellowship (LU 2336/2-1) by the DFG. HP is supported by the German Federal Ministry of Education and Research and Horizon2020 through the European Joint Programme on Rare Diseases ‘GENOMIT’ (01GM1920A). The Genotype-Tissue Expression (GTEx) Project was supported by the Common Fund of the Office of the Director of the National Institutes of Health, and by NCI, NHGRI, NHLBI, NIDA, NIMH, and NINDS. The authors gratefully acknowledge all participants in GTEx, including donors and previous analysts, who made this work possible. The authors also thank Yuanhua Huang, Marc Jan Bonder, Moritz von Scheidt, Hansi Weissensteiner, Bastian Linder and Andrew Dahl for giving us valuable feedback and analysis support.

## Ethical approval

This research was conducted under the ethical approval from dbGAP for access to individual level whole-genome sequencing (WGS) and RNA sequencing data in GTEx version 8^32^ (Accession: phs000424.v8.p2), through controlled access application number 23740.

## Author contributions

SW, MH and NC wrote the paper. NC designed the study. MH supported the mediation analysis. XG supported the data pre-processing and cell-type interaction analyses. PeK and FPC supported the association model testing. PaK and LSL performed the mtscATACseq analysis. HP supported the transcript processing analysis. SW performed all analyses. NC and MH supervised the work. All authors reviewed the paper.

## Data availability

Individual level sequencing data on GTEx samples used in this study are from dbGAP (Accession: phs000424.v8.p2) under controlled access application number 23740. We provide summary statistics on all significant findings in **Supplementary Tables**.

## Code availability

Publicly available tools that are used in data analyses are described wherever relevant in **Methods**. Custom code for performing all analyses in the paper is available at https://github.com/BioBuild/mtDNA-heteroplasmy.

## Extended Data Figures

**Extended Data Figure 1:**
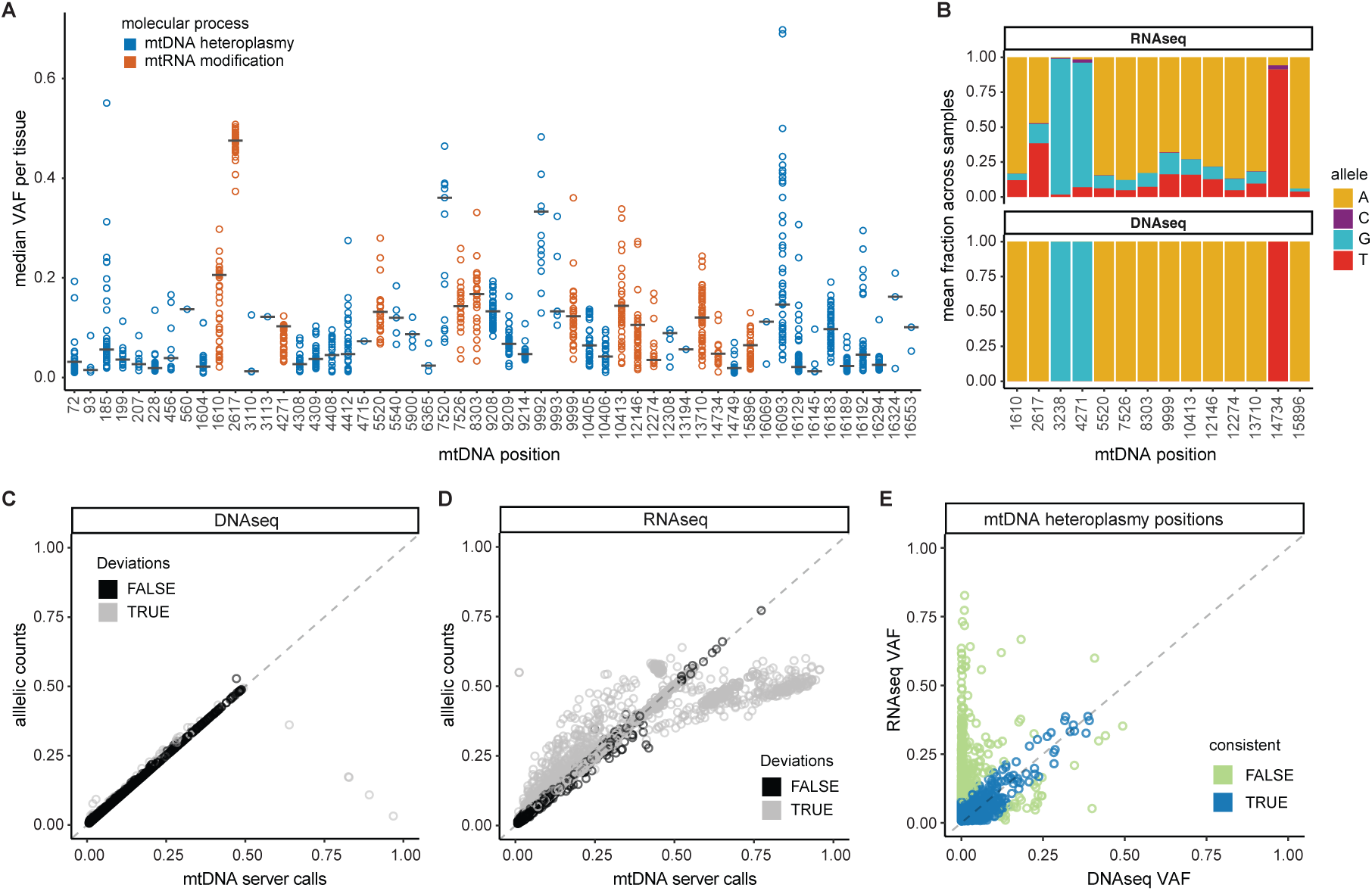
**A.** Median apparent heteroplasmy VAF per tissue at the top 10 % most variable apparent mtDNA heteroplasmy positions; each circle is a tissue. Grey bars indicate the median across tissues, colours indicate if the position is an RNA modification site. **B.** Mean of the fraction of alleles across samples detected in RNAseq and WGS at mtRNA modification sites in GTEx Whole Blood; colours indicate the respective alleles. **C.** Correlation VAF returned by the variant caller mtDNA server and the fraction of allelic counts obtained from WGS in GTEx Whole Blood; each dot is an apparent heteroplasmic site; colours indicate if the sum of MajorLevel and MinorLevel returned by mtDNA server deviated from 1.00 by >= 10 median absolute deviations (MADs). **D**. Correlation VAF returned by the variant caller mtDNAserver and the fraction of allelic counts obtained from RNAseq in GTEx Whole Blood; each dot is an apparent heteroplasmic site;; colours indicate if the sum of MajorLevel and MinorLevel returned by mtDNA server deviated from 1.00 by >= 10 median absolute deviations (MADs). **E.** Correlation of VAF quantified from RNAseq and from WGS data at mtDNA heteroplasmic sites in GTEx Whole Blood; consistency between heteroplasmy levels found in both data types is defined as <= 3 MADs.

**Extended Data Figure 2:**
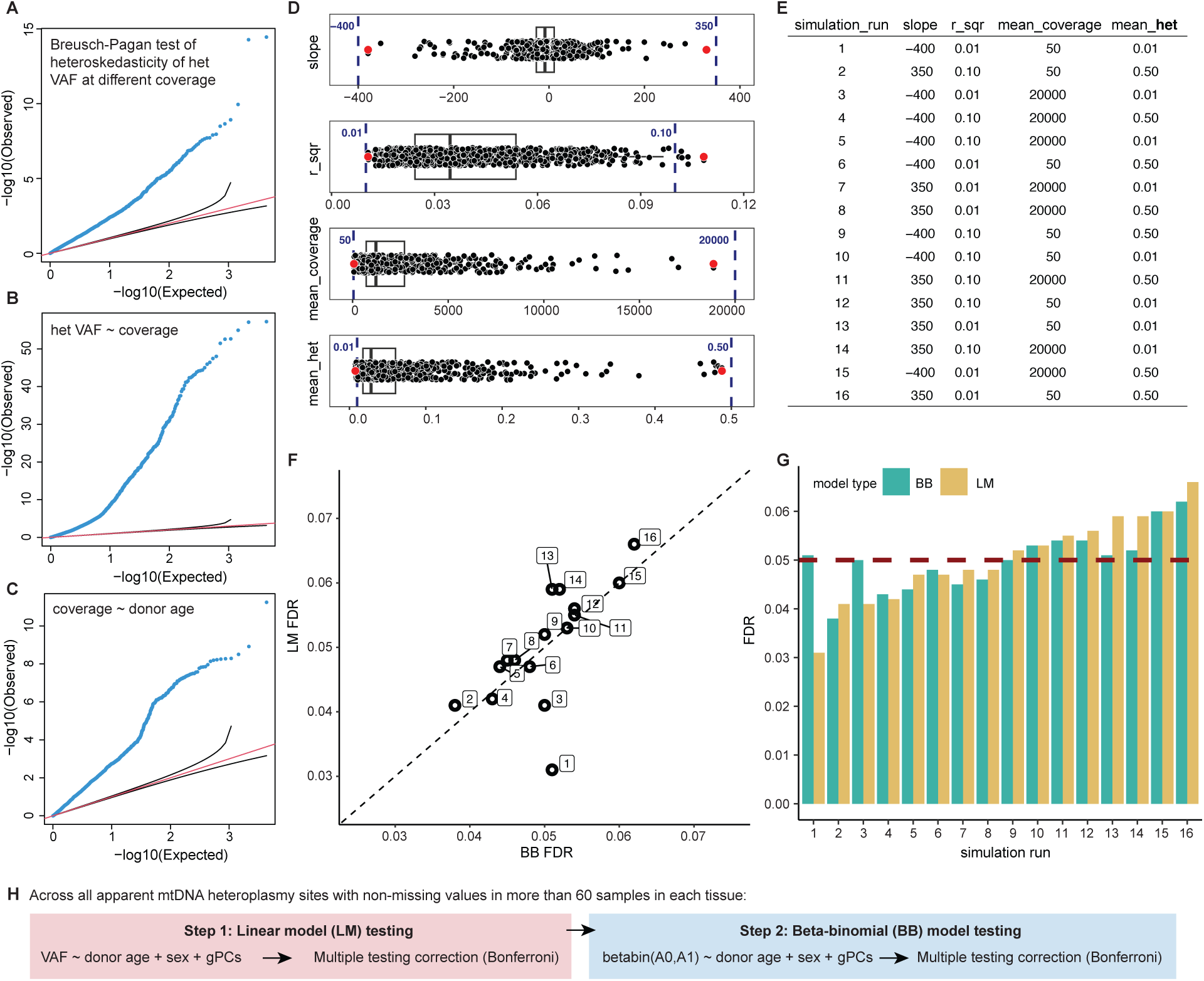
**A.** QQplot for Breusch-Pagan test for heteroskedasticity in apparent heteroplasmy VAF at different RNAseq coverage. **B.** QQplot for association results between heteroplasmy VAF and site-specific RNAseq coverage across all tissues. **C.** QQplot for association results between RNAseq coverage at all apparent mtDNA heteroplasmic sites and donor age across all tissues. For all QQplots in **A-C**, dots show results at all apparent heteroplasmic sites across all tissues; red line indicates y=x, black ranges show 95% confidence interval for null results. **D.** Boxplots of real data ranges identified in GTEx v8 across all apparent mtDNA heteroplasmy in all tissues for: regression coefficient between donor age and RNAseq coverage (slope); r2 values between donor age and RNAseq coverage (r_sqr); mean RNAseq coverage (mean_coverage) and mean apparent mtDNA heteroplasmy VAF (mean_het); red dots indicate minimum and maximum values identified in all tissues and apparent mtDNA heteroplasmic sites across GTEx v8; blue dashed lines and associated values indicate the range of values we use in our simulations; box shows interquartile range (IQR); median value is shown by the straight line in the box; whiskers show 1.5 x IQR. **E.** Table showing the 16 simulation runs with different parameters derived from realistic ranges across GTEx tissues; each simulation assumes no relationship between heteroplasmy VAF and donor age. **F**. Plot comparing the false discovery rate (FDR) for LM and BB models testing for relationships between heteroplasmy and donor age in synthetic data to the theoretical uniform distribution for LM and BB models; simulation runs are indicated by their numbers as shown in E. **G:** Barplot showing the estimated FDR for LM and BB models for each simulation run. **H.** Schematic showing our recommended two-step association testing approach using a combination of LM and BB models.

**Extended Data Figure 3:**
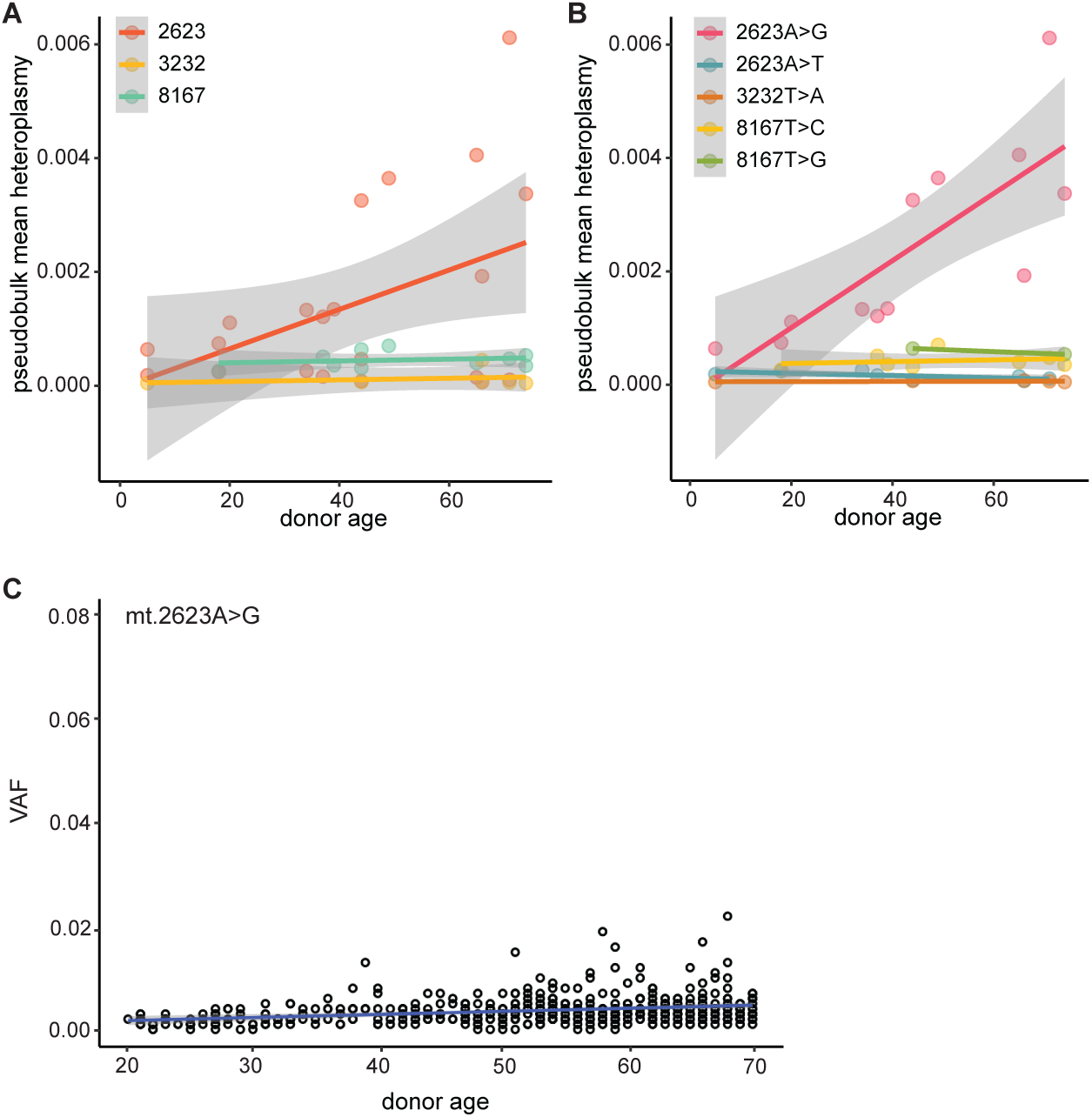
**A.** mtscATAC-seq pseudobulk mean heteroplasmy at position mt.2623, mt.3232, mt.8167 in human PBMCs across donor age. **B.** mtscATAC-seq pseudobulk mean heteroplasmy for shared variants detected at position mt.2623, mt.3232, mt.8167 in human PBMCs across donor age. **C**. VAF at mt.2623A>G in GTEx v8 whole blood RNAseq data across donor age.

**Extended Data Figure 4:**
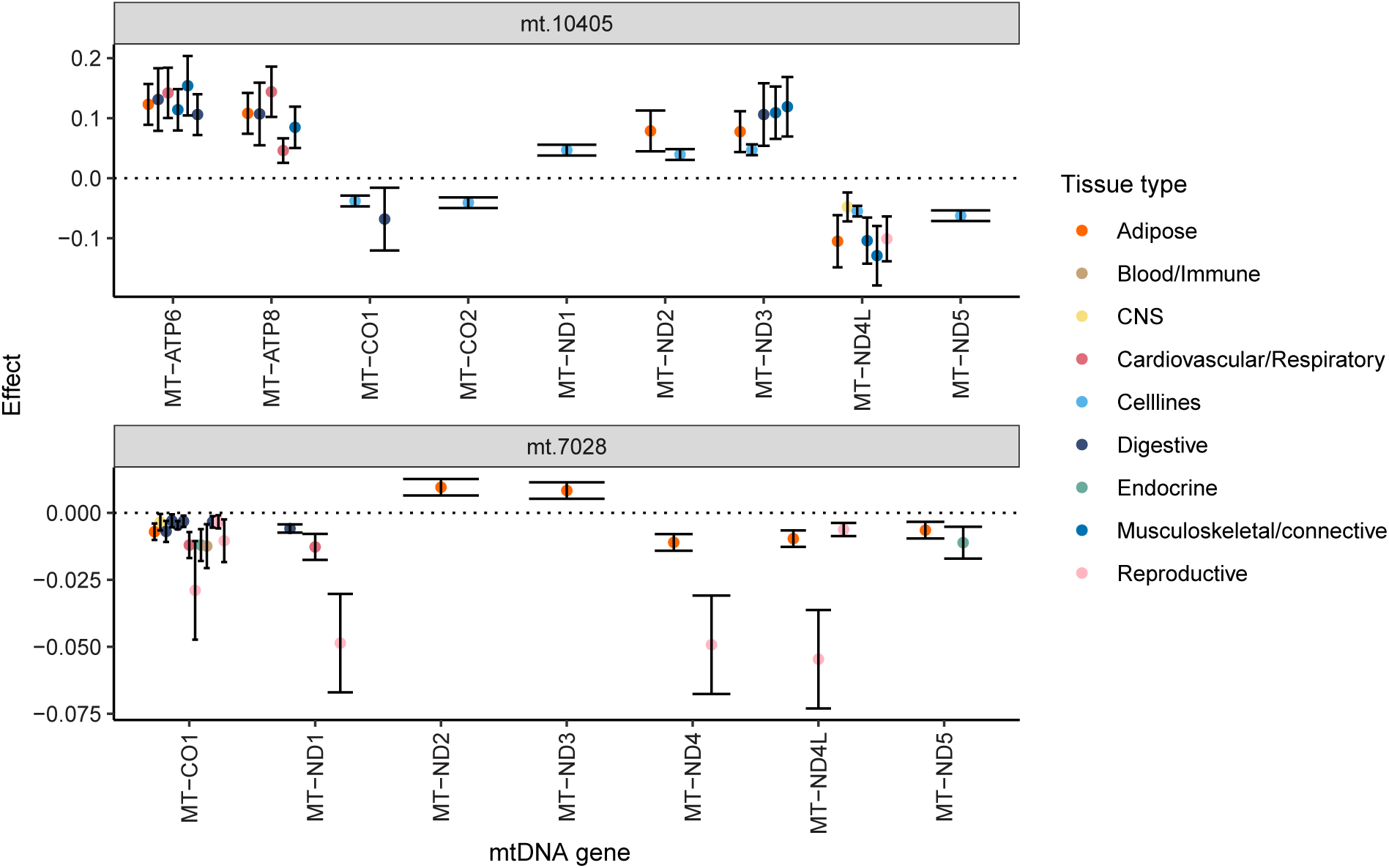
Plot showing the significant cis-eQTL effect sizes at mt.10405 (MT-TR) and mt.7082 (MT-CO1); each dot represents a significant cis-eQTL effect on a mtDNA gene in a tissue; dots are coloured by tissue type; error bars represent the 95% CI of the effect sizes.

**Extended Data Figure 5:**
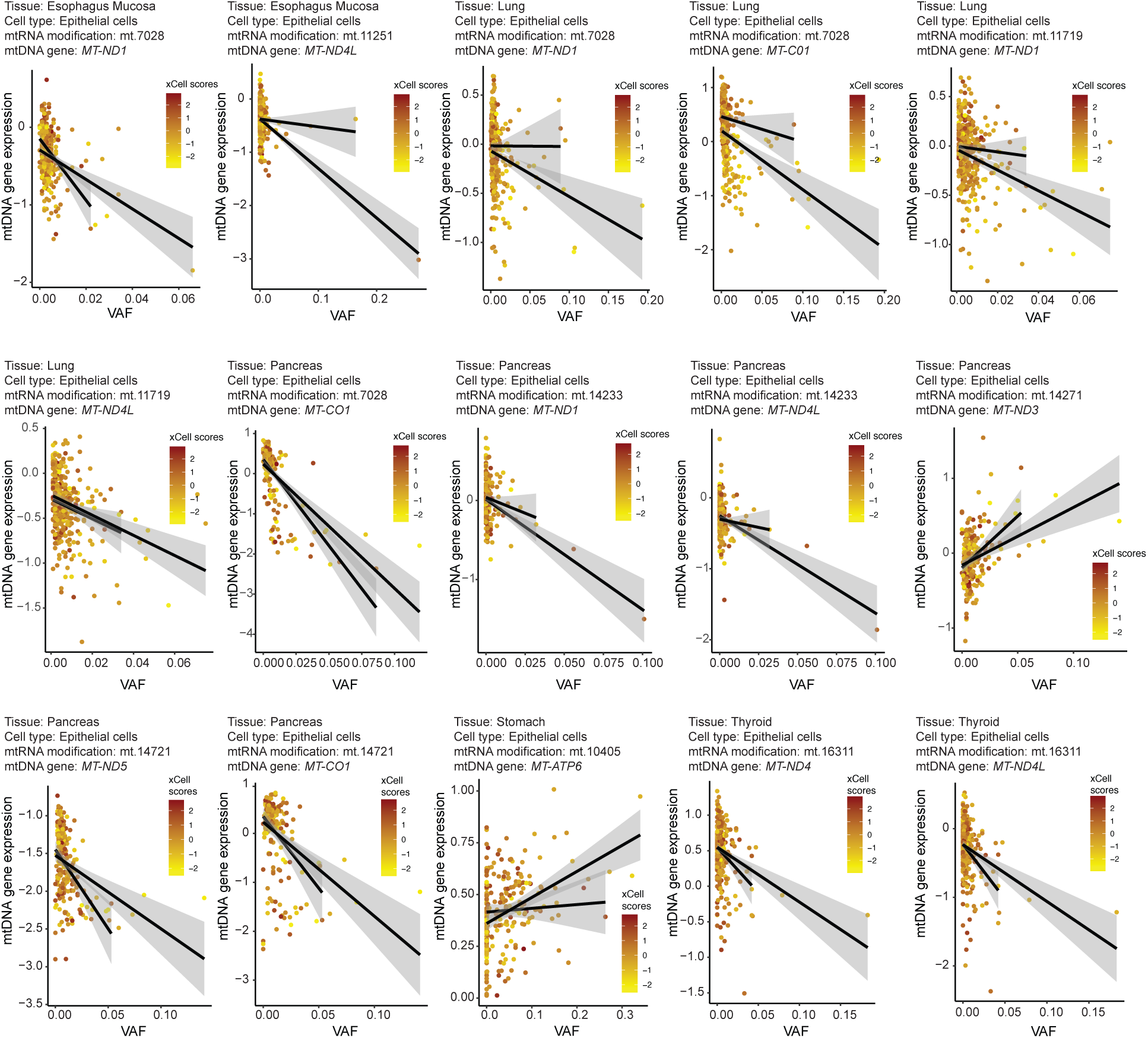
Significant interaction effects on tissue-specific mtDNA gene expression levels between VAFs at apparent mtDNA heteroplasmy and xCell scores of enriched cell types in each tissue^34^. For each significant interaction we show the VAF of the apparent mtDNA heteroplasmy on the x axis, the residuals of log(TPM+1) values for gene expression levels of mtDNA genes (corrected for PEER factors calculated per tissue) on the y axis, the xCell scores of the tested cell type in colour, and the black two lines show the linear relationships between the VAF and mtDNA gene expression levels at high (>mean) and low (<=mean) xCell scores, for visualization of the difference in effects at high and low xCell scores.

**Extended Data Figure 6:**
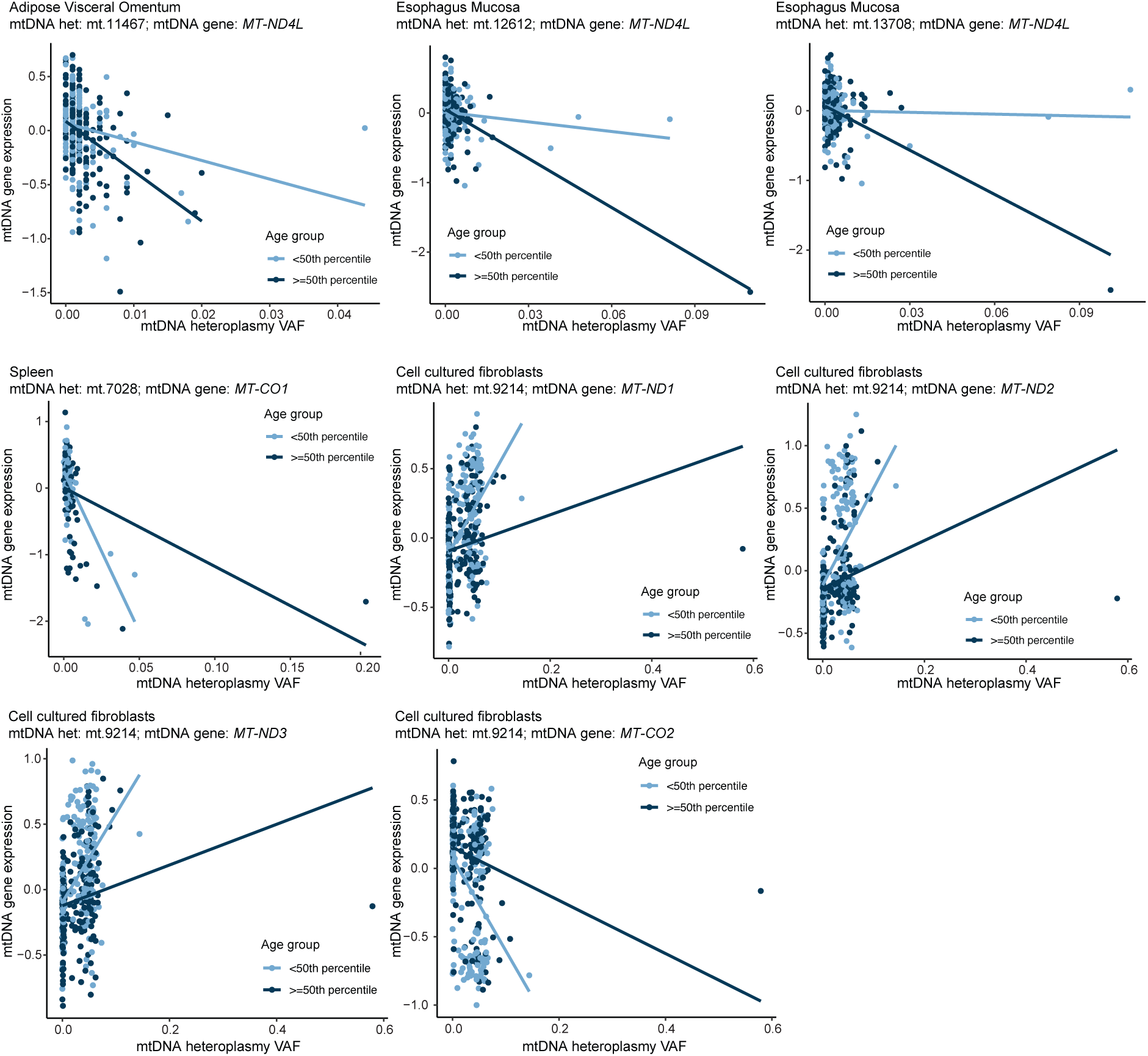
Significant interaction effects on tissue-specific mtDNA gene expression levels between VAFs at apparent mtDNA heteroplasmy and donor age. For each significant interaction we show the VAF of the apparent mtDNA heteroplasmy on the x axis, the residuals of log(TPM+1) values for gene expression levels of mtDNA genes (corrected for PEER factors calculated per tissue) on the y axis, the donor age groups (<50TH percentile and >= 50 percentile) in colour, along with lines of the respective colours showing the linear relationships between the VAF and mtDNA gene expression levels at high (>=50 percentile) and low (<50th percentile) donor ages, for visualization of the difference in effects at high and low donor ages.

