## SupplementaryMaterials for "Tissue-specific apparent mtDNA heteroplasmy and its relationship with ageing and mtDNA gene expression"

### Supplementary Methods

#### GTEx sample selection

We use nuclear DNA genotypes of GTEx samples to estimate their genetic ancestry and relatedness. We obtain 5,745,305 autosomal, biallelic SNPs in 838 GTEx samples from the variant call set from whole-genome sequencing (WGS) data in version 8[1] (Accession: phs000424.v8.p2, application #23740), having filtered the whole variant call set on minor allele frequency (MAF > 5%), p-value of violation of Hardy Weinberg Equilibrium (HWE > $10^{-6}$), and missingness (< 0.1). We then use KING[2] to identify related samples among all GTEx samples. Two pairs of individuals in GTEx are related up to third-degree (kinship >= 0.04419), though only marginally (kinship among pairs = 0.0477 and 0.0657), so we do not remove them from analyses.

To identify individuals of European ancestry, we select 180,936 Linkage disequilibrium-pruned (LD r2 < 0.2) common SNPs from the autosomes that overlap between GTEx and 1000 Genomes Project Phase 3 (1000G)[3] using PLINKv1.9[4], obtain their loadings of GTEx samples on principal components (PCs) identified in 1000G samples using the same SNPs, and project the GTEx samples onto the 1000G PCs with LDAK[5]. We select the 684 GTEx samples that cluster with the 1000G samples from the EUR superpopulation. We then build a genetic relatedness matrix (GRM) using 5,523,421 common SNPs (MAF > 5%, missingness < 0.1, P value for HWE > $10^{-6}$) in the European samples in GTEx, and derived PCs from this relatedness matrix and use them as covariates in all analyses we perform in this paper.

#### Quality control and custom filters for apparent mtDNA heteroplasmy calls

We perform stringent quality control on raw apparent mtDNA heteroplasmy calls from both DNA and RNAseq data. First, we only consider tissues with >= 60 samples. Second, we keep apparent mtDNA heteroplasmy calls identified at positions with adequate coverage (supporting either alleles; >= 10 reads on both forward and reverse strands), and which are observed in 5 samples or more in a tissue.

Third, we identify inconsistencies between variant calls returned by mtDNA server and allelic counts by applying a median absolute deviation (MAD) threshold of >= 10 between the two (**Supplementary Figure S4**). We find these inconsistencies are exclusively occurring in RNAseq variant calls, that is they are absent in DNAseq, and are driven by differences in RNAseq coverage mapping to the reverse and the forward sequencing strand (**Extended Data Figure 1, Supplementary Figure S2**). We conclude that these inconsistencies are reflecting strand bias not accounted for by mtDNAserver's internal QC procedure, which is developed in the context of mtDNAseq variant calling. We think it is likely this systematic strand bias is occuring in the maximum likelihood step, which is applied independently to each strand[6]. To remedy this, we quantify the strand bias as $\Delta_{STRAND} = abs(C_{REV} - C_{FWD})/C_{TOT}$, where $C_{REV}$ := coverage on reverse strand; $C_{FWD}$ := coverage on forward strand, $C_{TOT}$ := total coverage on both strands. We remove apparent heteroplasmy variant calls exceeding $\Delta_{STRAND} > 0.5$ from further analysis (**Supplementary Figure S4**).

The Spearman correlation r2 between heteroplasmy levels obtained from variant calling and allelic counts in mtDNA heteroplasmy and mtRNA modifications before this filter are 0.945 and 0.92 respectively, and after this filter they are 0.98 and 0.92 respectively. The remaining inconsistency between heteroplasmy levels obtained from variant calling and allelic counts specific to RNA modifications (r2 = 0.92) is accounted for by substituting heteroplasmy calls with allelic counts for RNA modifications throughout.

Fourth, we observe inconsistencies between heteroplasmy levels called from WGS compared with those called from RNAseq, which we quantify as $\Delta_{LEVELS} = abs(VAF_{RNAseq} - VAF_{DNAseq})$, where $VAF_{RNAseq}$ := heteroplasmy calls RNAseq; $VAF_{DNAseq}$ := heteroplasmy calls DNAseq; $\Delta_{LEVELS}$ inconsistent >= 3 MADs (**Extended Data Figure 1**). Since inconsistencies in $\Delta_{LEVELS}$ cannot be attributed to any particular transcript type or region of the mtDNA genome, but instead are found to be driven by the low coverage segment in RNAseq (total coverage after strand bias correction; **Supplementary Figure S3**), we assess the potential confounding due to this effect in contrast to actual RNAseq specific differences during model selection for association testing.

#### Testing for tissue enrichment

To test for CNS enrichment of mtRNA modifications, we group GTEx tissues into broader tissue categories and test CNS vs. the remaining tissues[7]. Since different numbers of tissues and multiple tissues from the same donors have been sampled we apply a downsampling procedure to disentangle these effects when performing CNS enrichment testing. Specifically, we group RNA modification sites by the nDNA encoded enzymes that catalyse the respective RNA methylations (TRMT10C[8–11], TRMT61B[12], unknown). We subsample a set of 10 tissues per donor, contrast the number of methylated sites in CNS tissues vs. the remaining tissues, and apply Fisher's exact test to obtain odds ratios as the enrichment effect. We repeat this sampling-testing procedure 1000 times and calculate the mean and 95 % confidence intervals (CI) of the odds ratios to assess the significance of the m1A/G tRNA methylation CNS enrichment (CI = 1.96*sd(OR), **Figure 2G**).

#### Assessing potential confounders in heteroplasmy estimates

To estimate the extent of heteroskedasticity in heteroplasmy levels across the observed range of RNA-seq coverage values, we fit a linear model VAF ~ C, where C denotes RNA-seq coverage (**Extended Data Figure 2A)**. To assess whether heteroskedasticity would be expected in VAF calls at different RNAseq coverage values, we apply the studentized Breusch-Pagan test, as implemented in the lmtest R package (v0.9-40) on the residuals of the VAF ~ C model described above. Next, to assess whether there is a relationship between RNAseq coverage and donor age, we applied the model C ~ A, where A represents donor age (**Extended Data Figure 2C)**. To account for multiple testing across all analyses, we apply a study-wide Bonferroni correction. Finally, we use the above tests to obtain parameters for simulating a realistic range of heteroskedasticity in apparent heteroplasmic and their relationships with RNAseq coverage and donor age in the GTEx cohort, as follows (**Extended Data Figure 2D**): **slope** := slope values for each position from the linear model C ~ A; **r_sqr** := R2 values for each position from the linear model C ~ A; **mean_coverage** := mean coverage values for each position; **mean_het** := mean VAF values for each position.

#### Simulations

To assess the calibration of linear models (LM) and beta-binomial (BB) models in testing for phenotype relationships in RNAseq derived heteroplasmy levels, where RNAseq coverage by itself and in correlation with the phenotype of interest may induce spurious relationships due to pervasive heteroskedasticity, we perform the following simulations using rounded values of the parameters observed in the GTEx data as described above: first, we simulate mean read coverage **mean_coverage** that varied linearly with donor age with effect **slope**, where ranges of both mean_coverage and slope tested are derived from the highest and lowest real measures of these parameters in GTEx RNAseq data (**Supplementary Table S3**). The intercept in the model is linked to the mean coverage. We then draw reads supporting the heteroplasmic allele from a BB distribution parameterised by **mean_beta** and **var_beta,** the ranges of which are derived from the highest and lowest real measures of mean and variance of heteroplasmy levels in GTEx RNAseq data (**Supplementary Table S3**). Importantly, in our simulation, these **mean_beta** and **var_beta** values are not related to donor age. In total, we set up 16 simulations of 500 donors and 1000 heteroplasmic variations each, with different combinations of values for **mean_coverage**, **slope**, **mean_beta** and **var_beta** (**Supplementary Table S7**). Finally, we test for association between the simulated heteroplasmy levels and donor age using both BB and LM, and count the rate of false positive (FP) findings.

#### mtscATACseq data processing and analysis

Peripheral blood mononuclear cells (PBMCs) from healthy donors were processed via mitochondrial single-cell ATAC-seq (mtscATAC-seq) as previously described[13]. Following sequencing via the Illumina NovaSeq platforms, fastq-files were processed with cellranger-atac (version 2.1.0)[14] using a NUMT blacklisted hg38 reference genome. Mitochondrial DNA (mtDNA) variant calling was performed using mgatk (version 0.7.0)[15]. High-confidence somatic mtDNA variants were identified using the following filtering parameters: variance-mean ratio >=0.01, coverage >5, strand correlation >0.65, and number of cells with at least two counts of the variant in both strands >=1. For the indicated positions/variants a linear regression of the pseudo bulk mean heteroplasmy over donor age was performed.

#### Mediation analysis between apparent mtDNA heteroplasmy, donor age and mtDNA gene expression

For the 10 apparent mtDNA heteroplasmies for which we find significant associations with donor age and with mtDNA gene expression in the same tissues, we further investigate the causal relationships between them (see graphical summary in **Figure 5A**).

Specifically, to establish a mediation relationship between a given apparent mtDNA heteroplasmy (VAF), mtDNA gene expression levels (GE), and a donor's age (AGE) we require the following (covariates: donor sex, gPC1-5):

1) The apparent mtDNA heteroplasmy is significantly associated with donor age and this association remains when adjusting for mtDNA gene expression. We evaluate the Pearson correlation of the residuals from the following models to test this relationship, and consider this requirement fulfilled if the correlation is **significant**.

   (i) $\text{VAF} \sim \beta 0 + \beta 1 \cdot \text{AGE} + \beta 2 \cdot \text{covariates} + \varepsilon$
   (ii) $\text{GE} \sim \beta 0 + \beta 1 \cdot \text{AGE} + \beta 2 \cdot \text{covariates} + \varepsilon$

2) The apparent mtDNA heteroplasmy is significantly associated with mtDNA gene expression, and this association remains when adjusting for donor age. We evaluate the Pearson correlation of the residuals from the following models to test this relationship, and consider this requirement fulfilled if the correlation is **significant**.
   (i) VAF ~ $\beta 0 + \beta 1 \cdot$ GE + $\beta 2 \cdot$ covariates + $\varepsilon$
   (ii) AGE ~ $\beta 0 + \beta 1 \cdot$ GE + $\beta 2 \cdot$ covariates + $\varepsilon$

3) Donor age is significantly associated with mtDNA gene expression, but this association disappears when adjusting for the apparent mtDNA heteroplasmy level. We evaluate the Pearson correlation of the residuals from the following models to test this relationship, and consider this requirement fulfilled if the correlation is **significant**.

   (i) AGE ~ $\beta 0 + \beta 1 \cdot$VAF + $\beta C \cdot$ covariates + $\varepsilon$
   (ii) GE ~ $\beta 0 + \beta 1 \cdot$ VAF + $\beta C \cdot$ covariates + $\varepsilon$

Multiple testing correction is conducted across all VAF/GE/AGE trios identified in all tissues (10281 instances), giving a significance threshold of $P < 0.05/10281$ (study-wide Bonferroni).

We interpret the outcomes of 1), 2) and 3) to establish mediation effects as follows (summarised in **Supplementary Table S12**):

AGE → VAF → GE: 1) = significant, 2) = significant, 3) = insignificant. We find 9/10 such instances.

AGE → GE → VAF: 1) = significant, 2) = insignificant, 3) = significant. We find 0/10 such instances.

For all other combinations of 1), 2), and 3), accounting for only 1/10 of the instances in our analysis, no conclusive interpretations can be drawn. This may be due to limited power in the dataset we use; future work on larger datasets with higher statistical power to identify subtle relationships in gene expression regulation may be able to uncover more interpretable relationships.

**Supplementary Figures**

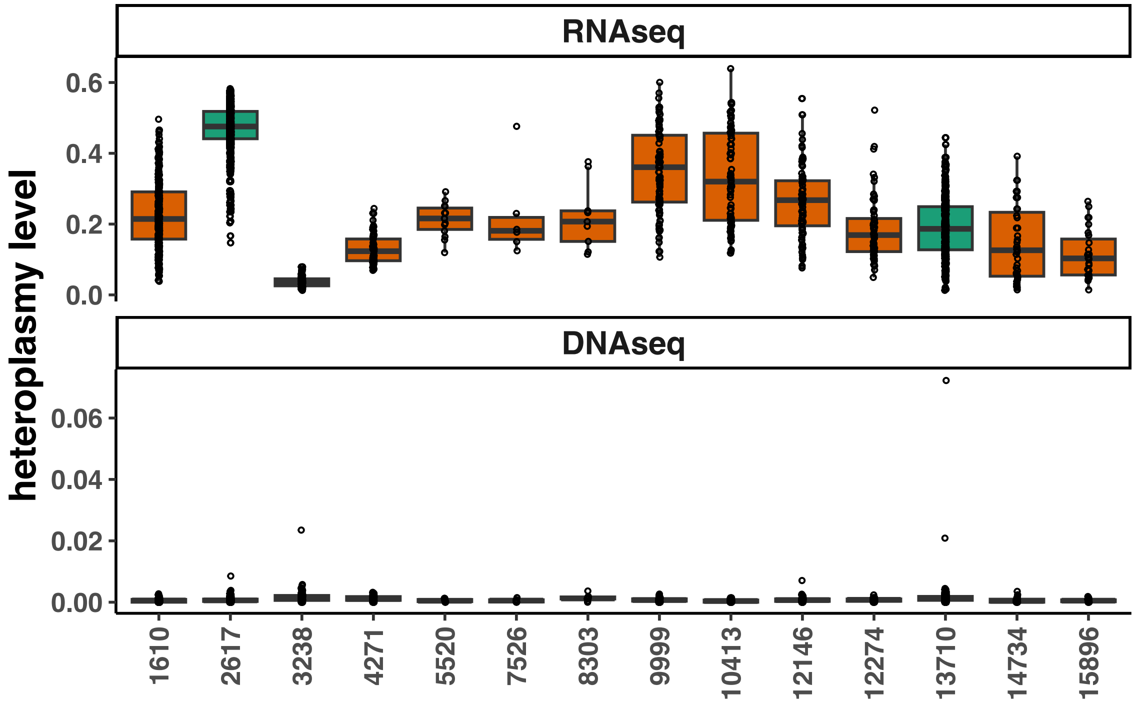

**Supplementary Figure S1:** VAFs at apparent heteroplasmy previously reported as being reflective of m1A/G RNA modification levels in 14 sites found in 548 donors in GTEx Whole Blood. **RNAseq:** VAFs detected from RNAseq. **DNAseq**: VAF detected from DNAseq.

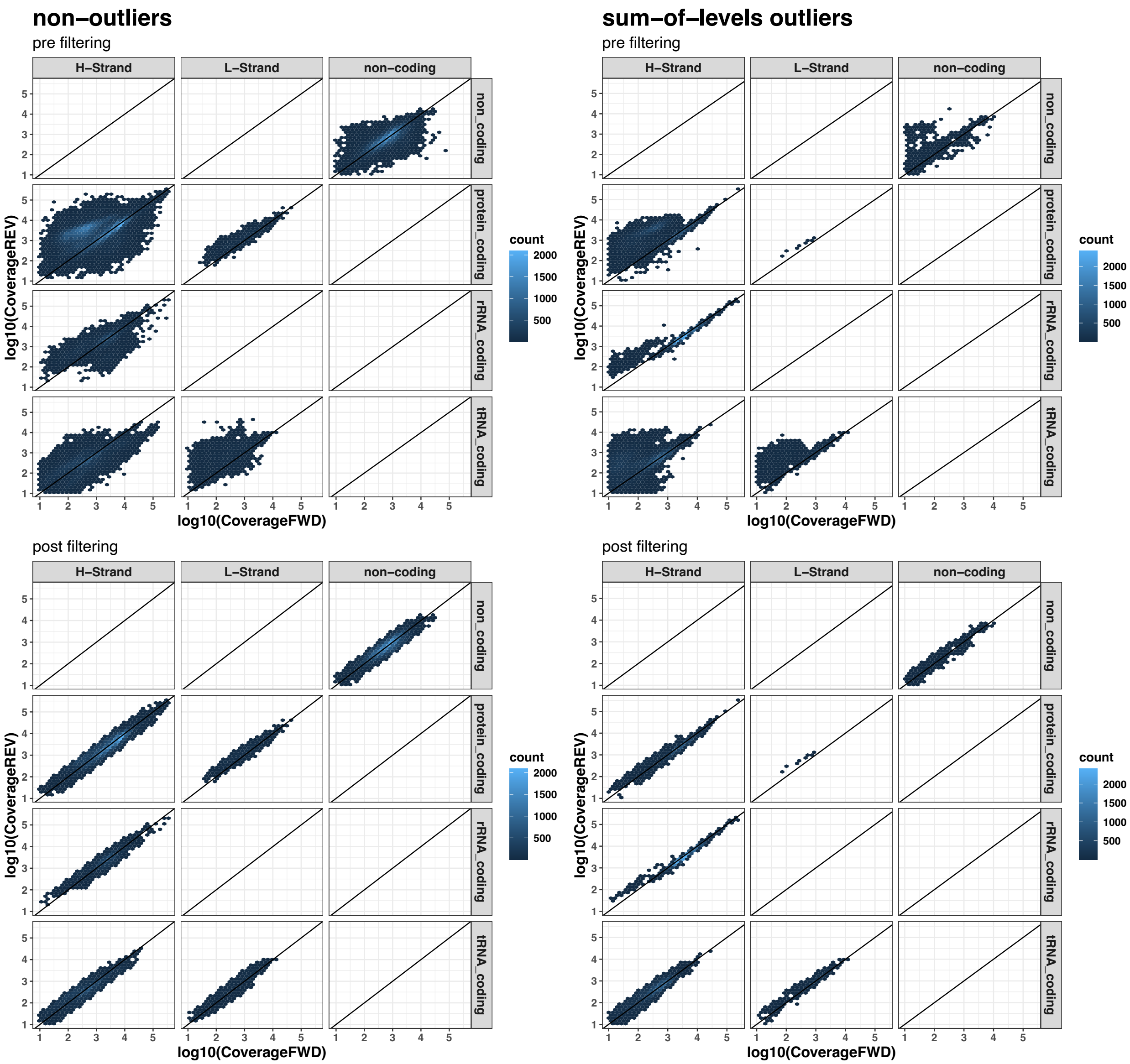

**Supplementary Figure S2:** Density estimates of log10 RNAseq coverage detected at the forward strand (FWD) vs. reverse strand (REV). Facetting indicates the feature strand (H-strand, L-strand or non-coding) along the x-axis and the transcript type of the features overlapping the respective heteroplasmy along the y-axis. sum-of-level-outliers := MAD of the total sum of VAFs at a given heteroplasmy >= 10. strand bias threshold := delta_frac >= 0.5 (delta_frac = abs(CoverageREV - CoverageFWD)/Coverage)). **Top left:** heteroplasmy which does not show sum-of-level-outlier VAFs before filtering for coverage strand bias. **Top right:** heteroplasmy which does show sum-of-level-outlier VAFs before filtering for coverage strand bias. **Bottom left:** heteroplasmy which do not show sum-of-level-outlier VAFs after filtering for coverage strand bias. **Bottom right:** heteroplasmy which do show sum-of-level-outlier VAFs after filtering for coverage strand bias.

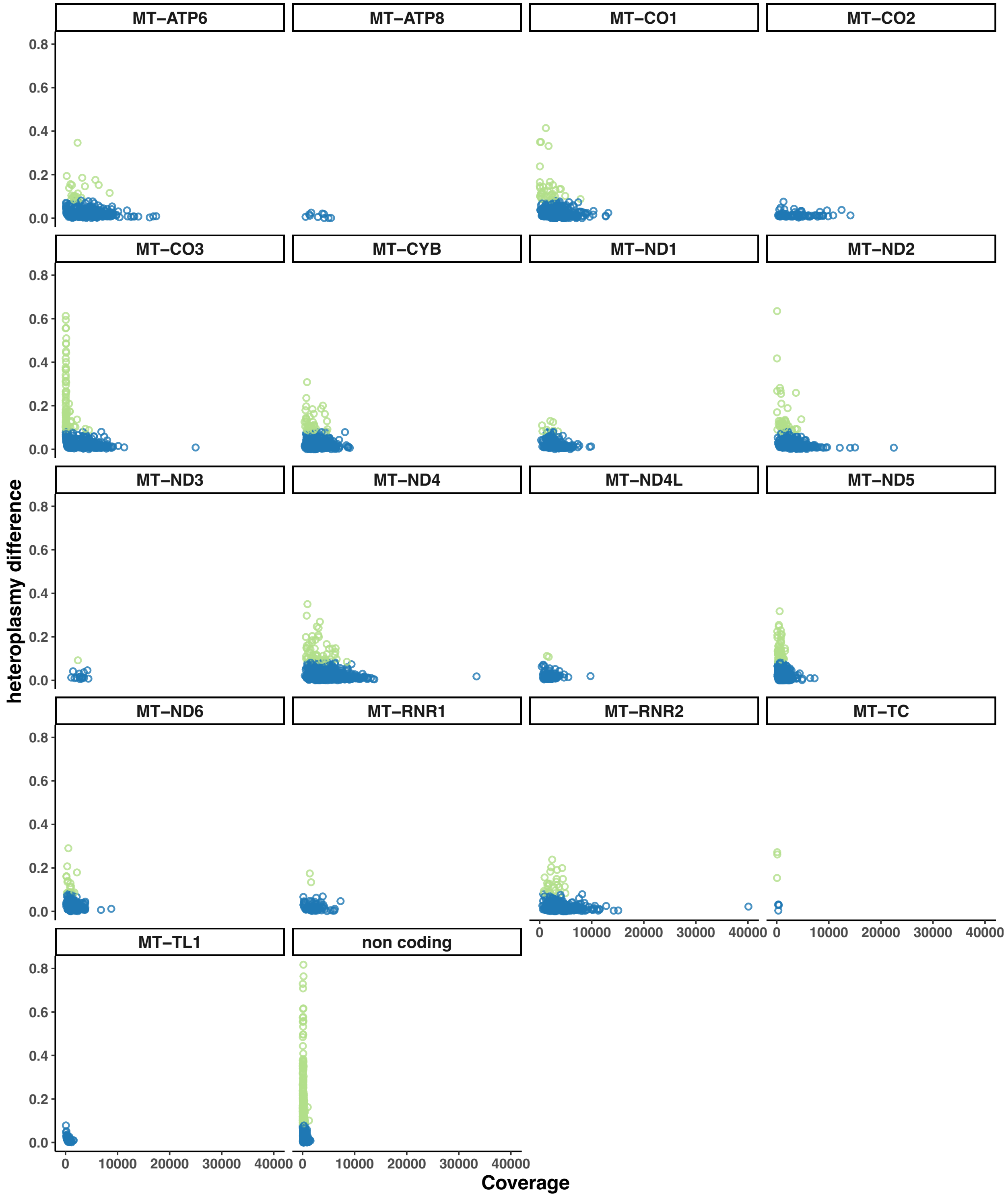

**Supplementary Figure S3:** RNAseq coverage at apparent heteroplasmy detected from RNAseq in GTEx Whole Blood vs. the absolute difference in VAFs detected from RNAseq vs. DNAseq. VAFs at apparent heteroplasmy consistent between the two data types were defined as >= 3 MAD of heteroplasmy difference. Consistent VAFs are highlighted in blue, inconsistent ones in green. Each facet indicates the mtDNA encoded gene which overlaps the respective apparent heteroplasmy.

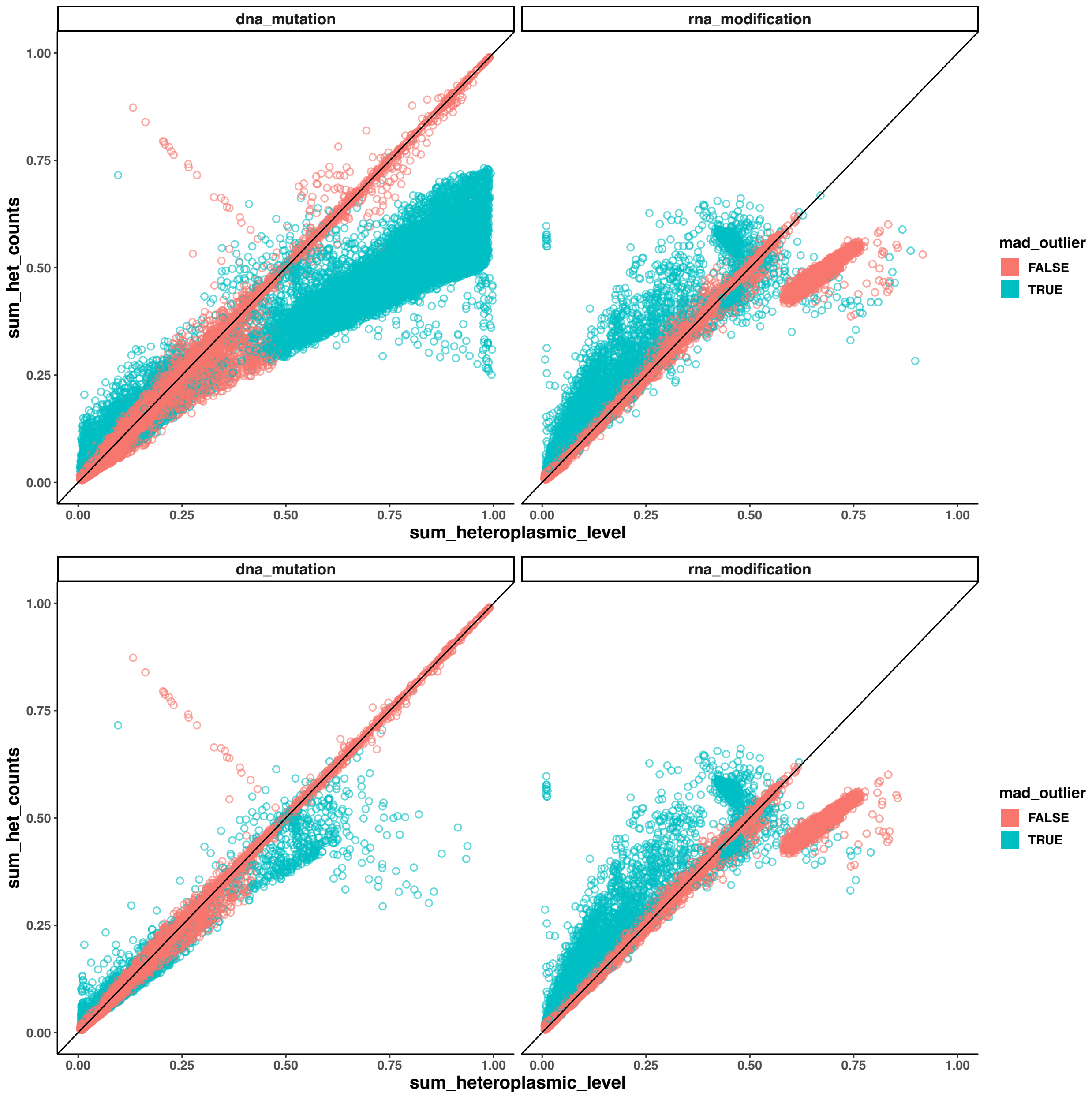

**Supplementary Figure S4:** Sum of VAFs detected per apparent heteroplasmy by mtDNA server across all RNAseq samples (sum_heteroplasmic_level) vs. the sum of allelic counts (sum_het_counts). Colours indicate if the sum of MajorLevel and MinorLevel returned by mtDNA server deviated from 1.00 by >= 10 MADs. **Upper left:** VAFs at mtDNA heteroplasmy pre custom strand-bias filtering. **Upper right:** VAFs at mtRNA modifications, pre custom strand-bias filtering. **Lower left:** VAFs at mtDNA heteroplasmy after custom strand-bias filtering. **Lower right:** VAFs at mtRNA modifications after custom strand-bias filtering. Custom strand bias filter was defined as 10 MADs <= of the absolute fraction of the differences in RNAseq coverage detected on the reverse (REV) and the forward strand (FWD) ($\Delta_{STRAND}$ >= 0.5; $\Delta_{STRAND}$ = abs($C_{REV}$ - $C_{FWD}$)/$C_{TOT}$, where $C_{REV}$ := coverage on reverse strand; $C_{FWD}$ := coverage on forward strand, $C_{TOT}$ := total coverage on both strands.)

**Supplementary Table Legends**

**Supplementary Table S1**: Number of apparent heteroplasmy across all individuals and tissues (n_heteroplasmy), and number of unique mtDNA positions (n_positions), number of tissue-specific samples (n_samples) and number of donors (n_donors) these apparent heteroplasmy are identified in, remaining after each filtering step (step, filter_type) in the quality control pipeline.

**Supplementary Table S2**: Summary of the total number of apparent heteroplasmy (n_total_apparent_heteroplasmy) and the breakdown of the number of mtDNA heteroplasmy (n_mtDNA_heteroplasmy) and mtRNA modifications (n_mtRNA_modification) found per tissue (tissue, annot) in the total number of samples (n_samples) after the whole quality control pipeline.

**Supplementary Table S3**: Table of all known mtRNA modification sites in one protein coding transcript *MT-ND5*, one mt-rRNA *MT-RNR2*, and 22 mt-tRNAs; for each modification site we show the mtDNA gene and its properties (gene_name, strand_name, gene_start, gene_end), the mtDNA and transcript positions of the mtRNA modification (genomic_pos, transcript_pos), the nature of the modification (rna_modification); for each mt-tRNA modification we further show the amino acid it translates (encoded_aa), the start and end mtDNA position of the anticodon (anticodon_start, anticodon_end), the mt-tRNA and mtDNA trinucleotide sequence of the anticodon (anticodon_rna, anticodon_dna), and whether there is a disease association with an anticodon change at this position in MITOMAP[16].

**Supplementary Table S4**: Significant association results between apparent heteroplasmy VAF and site-specific RNAseq coverage at 2086 out of 4334 apparent heteroplasmy positions tested; for each significant association we show the tissue it is tested in (tissue, annot), the apparent heteroplasmy position (het_id), the effect of RNAseq coverage on apparent heteroplasmy VAF and its standard error (effect, se), the raw and the study-wide Bonferroni adjusted p value of the association (p_value, adj_p_value).

**Supplementary Table S5**: Significant Breusch-Pagan test results for heteroskedasticity in apparent heteroplasmy VAF at different RNAseq coverage at 70 out of 4334 apparent heteroplasmy positions tested; for each significant result we show the tissue it is tested in (tissue, annot), the apparent heteroplasmy position (het_id), the Breusch-Pagan test statistic (bptest_statistic), and the raw and the study-wide Bonferroni adjusted p value of the association (bptest_p_value, bptest_adj_p_value).

**Supplementary Table S6**: Significant association results between RNAseq coverage at apparent mtDNA heteroplasmic positions and donor age at 830 out of 4,334 apparent mtDNA heteroplasmy positions across tissues; for each significant association we show the tissue it is tested in (tissue, annot), the apparent heteroplasmy position (het_id), the effect of donor age on RNAseq coverage and its standard error (effect, se), the raw and the study-wide Bonferroni adjusted p value of the association (p_value, adj_p_value).

**Supplementary Table S7**: The minimum and maximum values of the following parameters (extreme_value_type) on the relationship between donor age and RNAseq coverage at apparent heteroplasmy positions with these values, derived from samples across tissues in GTEx at apparent heteroplasmy positions (tissue, Pos): regression coefficient between donor age and RNAseq coverage (slope), r2 values between donor age and RNAseq coverage (r_sqr), p value of association between donor age and RNAseq coverage (p_value), as well as the mean RNAseq coverage (mean_coverage) and the mean and variance of the apparent heteroplasmy VAF (mean_het, var_het) at these positions.

**Supplementary Table S8**: Significant association results between apparent heteroplasmy VAF and donor age at 109 out of 4,334 apparent mtDNA heteroplasmy positions across tissues; for each significant association we show the tissue it is tested in (tissue, annot), the apparent heteroplasmy position and the mtDNA gene it is in (het_id, gene_name), the type of molecular process the apparent heteroplasmy is a result of (molecular_process), the putative type of mtDNA heteroplasmy where applicable (mtdna_het_type), the inherited and heteroplasmic alleles identified in the samples in GTEx at this position (inherited_allele, heteroplasmic_allele), the mean and standard deviation of the VAF at this position (mean_VAF, sd_VAF), and the number of samples in the association test; for each test we show the linear model effect size, standard error and raw and study-wide Bonferroni adjusted p values (LM_estimate, LM_std_error, LM_A_pval, LM_A_bonferroni), as well as the beta-binomial model effect size, standard error, raw and study-wide Bonferroni adjusted analytical p values (BB_estimate, BB_std_error, BB_A_pval, BB_A_bonferroni), and raw and study-wide Bonferroni adjusted empirical p values from permutations (BB_E_pval, BB_E_bonferroni).

**Supplementary Table S9**: Significant association results between apparent heteroplasmy VAF and mtDNA gene expression at 784 out of 4,334 apparent mtDNA heteroplasmy positions across tissues; for each significant association we show the tissue it is tested in (tissue, annot), the apparent heteroplasmy position and the mtDNA gene it is in (het_id, gene_name), the type of molecular process the apparent heteroplasmy is a result of (molecular_process), the putative type of mtDNA heteroplasmy where applicable (mtdna_het_type), the inherited and heteroplasmic alleles identified in the samples in GTEx at this position (inherited_allele, heteroplasmic_allele), the mean and standard deviation of the VAF at this position (mean_VAF, sd_VAF), the eQTL mtDNA gene (eqrl_gene_name, ensembl_id), and the number of samples in the association test; for each test we show the linear model effect size, standard error and raw and study-wide Bonferroni adjusted p values (LM_estimate, LM_std_error, LM_A_pval, LM_A_bonferroni), as well as the beta-binomial model effect size, standard error, raw and study-wide Bonferroni adjusted analytical p values (BB_estimate, BB_std_error, BB_A_pval, BB_A_bonferroni), and raw and study-wide Bonferroni adjusted empirical p values from permutations (BB_E_pval, BB_E_bonferroni).

**Supplementary Table S10**: Significant cell type interaction test results between apparent heteroplasmy VAF and mtDNA gene expression at 18 out of 784 significant apparent heteroplasmy eQTLs across tissues performed using a linear model; for each significant interaction we show the tissue it is tested in (tissue, annot), the apparent heteroplasmy position and the mtDNA gene it is in (het_id, gene_name), the type of molecular process the apparent heteroplasmy is a result of (molecular_process), the putative type of mtDNA heteroplasmy where applicable (mtdna_het_type), the inherited and heteroplasmic alleles identified in the samples in GTEx at this position (inherited_allele, heteroplasmic_allele), the mean and standard deviation of the VAF at this position (mean_VAF, sd_VAF), the eQTL mtDNA gene (eqrl_gene_name, ensembl_id), the cell type in which the interaction effect is found, and the number of samples in the interaction test; for each test we show the interaction effect size, standard error and raw and study-wide Bonferroni adjusted p values (interaction_estimate, interaction_std_error, interaction_pval, interaction_bonferroni).

**Supplementary Table S11**: 10 tissue-specific heteroplasmy positions that show both significant eQTL with mtDNA genes and donor age associations; for each position we show the tissue it is tested in (tissue, annot), the apparent heteroplasmy position and the mtDNA gene it is in (het_id, gene_name), the type of molecular process the apparent heteroplasmy is a result of (molecular_process), the putative type of mtDNA heteroplasmy where applicable (mtdna_het_type), the inherited and heteroplasmic alleles identified in the samples in GTEx at this position (inherited_allele, heteroplasmic_allele), the mean and standard deviation of the VAF at this position (mean_VAF, sd_VAF), the eQTL mtDNA gene (eqrl_gene_name, ensembl_id), and the number of samples in the association test; for each position we show the linear model effect size, standard error and raw and study-wide Bonferroni adjusted p values for both eQTL and donor age associations (test_LM_estimate, test_LM_std_error, test_LM_A_pval, test_LM_A_bonferroni); for all positions we determine whether the donor age effects on gene expression is likely mediated by heteroplasmy (med_via_het), whether the apparent heteroplasmy effects on gene expression is likely mediated by age (med_via_age), and whether the two tests are independent (independent), using mediation analyses shown in **Supplementary Table S12**.

**Supplementary Table S12**: Results of mediation analysis between apparent heteroplasmy, mtDNA gene expression and donor age using partial correlations at 10 tissue-specific heteroplasmy positions that show both significant eQTL with mtDNA genes and donor age associations; we show results of the following partial correlations (edge) corresponding to edges of the mediation shown on **Figure 5**: edge 1 (VAF ~ mtDNA gene expression | donor age), edge 2 (VAF ~ donor age | mtDNA gene expression) and edge 3 (mtDNA gene expression ~ donor age | VAF); for each edge we show the term we condition on (conditional_on), the Pearson correlation r (pearson_r), and the raw and Bonferroni adjusted p value of the correlation (pearson_p_value, pearson_adj_p_value).

**Supplementary Table S13**: Significant donor age interaction test results between apparent heteroplasmy VAF and mtDNA gene expression at 18 out of 784 significant apparent heteroplasmy eQTLs across tissues performed using a linear model; for each significant interaction we show the tissue it is tested in (tissue, annot), the apparent heteroplasmy position and the mtDNA gene it is in (het_id, gene_name), the type of molecular process the apparent heteroplasmy is a result of (molecular_process), the putative type of mtDNA heteroplasmy where applicable (mtdna_het_type), the inherited and heteroplasmic alleles identified in the samples in GTEx at this position (inherited_allele, heteroplasmic_allele), the mean and standard deviation of the VAF at this position (mean_VAF, sd_VAF), the eQTL mtDNA gene (eqrl_gene_name, ensembl_id), and the number of samples in the interaction test; for each test we show the linear model effect size, standard error and raw p values for both eQTL and donor age associations (test_LM_estimate, test_LM_std_error, test_LM_A_pval), as well as the interaction effect size, standard error and raw and study-wide Bonferroni adjusted p values (interaction_estimate, interaction_std_error, interaction_pval).

**Supplementary Table S14**: Replication of association analysis between VAF of mtRNA modifications with 5' gene expression at 7 p9 mt-tRNA modification sites with a protein coding 5' gene in a previous study using GTEx v6 data on Whole Blood[17], this time using data in GTEx v8 Whole Blood; for each association we show the tissue it is tested in (tissue, annot), the apparent heteroplasmy position and the mt-tRNA it is in (het_id, mt-trna), the  and the number of samples used in association testing, the 5' mtDNA gene (5prime_gene_name, ensembl_id), the effect of p9 mt-tRNA modification VAF on 5' mtDNA gene expression and p value identified in GTEx v6 in the previous study (v6_effect, v6_p_value), and the effect, standard error and p value of p9 mt-tRNA modification VAF on the same 5' mtDNA gene expression identified in this study using GTEx v8 Whole Blood (v8_effect, v8_std_error, v8_p_value).

**Supplementary Table S15**: Full results of the association analysis between VAF of mtRNA modifications with 5' gene expression at all identified p9 mt-tRNA modification sites with a protein coding 5' gene across tissues; for each association we show the tissue it is tested in (tissue, annot), the apparent heteroplasmy position and the mt-tRNA it is in (het_id, mt-trna), the  and the number of samples used in association testing, the 5' mtDNA gene (5prime_gene_name, ensembl_id), the effect, standard error and p value of p9 mt-tRNA modification VAF on the same 5' mtDNA gene expression identified in this study using GTEx v8 Whole Blood (VAF_effect, VAF_std_error, VAF_p_value), and whether this association is significant at the tissue or study-wide level using Bonferroni correct (VAF_sig_tissuewide, VAF_sig_studywide); we further show the results of a follow-up analysis testing if gene expression levels of these 5' mtDNA genes are associated with a 5' "cut" among p9 modified reads using a binomial regression (**Methods**): the effect of the 5' "cut", the standard error of the effect, the p value (cut_effect, cut_std_error, cut_p_value), and whether this association is significant at the tissue or study-wide level using Bonferroni correct (cut_sig_tissuewide, cut_sig_studywide).
